# The assembly of cancer-specific ribosomes by the lncRNA *LISRR* suppresses melanoma anti-tumour immunity

**DOI:** 10.1101/2023.01.06.523012

**Authors:** Sonia Cinque, Yvessa Verheyden, Sara Adnane, Alessandro Marino, Vicky Katopodi, Ewout Demesmaeker, Zorica Knezevic, Sarah Hanache, Roberto Vendramin, Alessandro Cuomo, Joanna Pozniak, Alvaro Cortes Calabuig, Marcella Baldewijns, Sébastien Tabruyn, Oliver Bechter, Maria Francesca Baietti, Elisabetta Groaz, Tiziana Bonaldi, Eleonora Leucci

**Affiliations:** Laboratory for RNA Cancer Biology, Department of Oncology, KU Leuven, Leuven, Belgium; Trace, Leuven Cancer Institute, KU Leuven, Leuven, Belgium; Department of Experimental Oncology, IEO, European Institute of Oncology IRCCS, Via Adamello 16, 20139 Milan, Italy; Laboratory for Molecular Cancer Biology, Center for Cancer Biology, VIB, Leuven, Belgium; Laboratory for Cytogenetics and Genome Research, Department of Human Genetics, KU Leuven, Leuven, Belgium; Department of Pathology, University Hospitals Leuven, 3000 Leuven, Belgium; TransCure bioServices, Archamps France; Department of General Medical Oncology UZ Leuven, Leuven, Belgium; Rega Institute for Medical Research, Medicinal Chemistry, KU Leuven, Leuven, Belgium; Department of Oncology and Hematology-Oncology, University of Milan, Milan, 20122 Italy

## Abstract

Although immune checkpoint blockade (ICB) has revolutionized cancer treatment, resistance mechanisms limit its clinical benefit. Here we characterise *LISRR*, a cancer-specific lncRNA highly expressed in melanoma patients refractory to ICB. In cells undergoing (therapeutic) stress, *LISRR* recruits DAZAP1 (Deleted in AZoospermia Associated Protein 1) to polysomes and drives the assembly of a subset of ribosomes at the endoplasmic reticulum, directing the synthesis of an immunosuppressive translatome. This includes the immune checkpoint PD-L1 and the enzymes necessary for building the glycocalyx, the sugar coat surrounding the cells. Notably, proper glycocalyx assembly is required for spermatozoa immune evasion during fertilization. Accordingly, targeting *LISRR* activates immune responses and re-sensitizes to ICB in co-culture models, *ex vivo* in patient explants, and *in vivo* in humanized patient-derived models. Our study reveals the contribution of lncRNAs to the generation of cancer-specific ribosomes and identifies an RNA-based cancer-specific strategy to overcome intrinsic resistance to ICB.

## Introduction

The introduction of immune checkpoint inhibitors has revolutionised cancer treatment, however intrinsic and acquired resistance mechanisms hamper the success of immune therapy in a significant portion of patients [1].

A key non-genetic mechanism fuelling the generation of Drug Tolerant Persister Cells (DTPCs) is the Integrated Stress Response (ISR), a cytoplasmic adaptive response to intracellular and extracellular stress stimuli (e.g., amino acids deprivation, inflammation, ER stress…), consisting of a global reduction of cap-dependent translation initiation followed by the activation of special transcriptional and translational programs orchestrated by ATF4. By coordinating the unfolded protein response, amino acid metabolism and oxidative stress responses, among the others [2, 3], the transcription factor ATF4 promotes DTPCs survival.

The selective recruitment of specific pro-survival mRNAs to ribosomes and their translation, is believed to rely mostly on structural motifs (e.g., G-quadruplexes, IRES…) on the mRNAs. Whether trans-acting factors are also involved in this recruitment process is yet unknown but is a likely and interesting possibility. Canonical coding and non-coding components of the translational machinery (e.g., ribosomal proteins, tRNAs, rRNAs) and their transcriptional, post-transcriptional and post-translational modifications [4] are emerging as critical regulators of translational rewiring in cancer. Whether long non-coding RNAs (lncRNAs), a large fraction of which is associated with ribosomes [5–8], also contribute to translation adaptation remains underexplored. It has been proposed that lncRNA binding to ribosomes may modulate their biogenesis and/or the recruitment/translation of single mRNAs [9–11]. Moreover a handful of these transcripts (∼2-7%) seems to produce (micro)peptides [12–15], some of which with a reported function [16–18]. The possibility that lncRNAs participate to the generation of cancer-specific ribosomes thus modifying the translational landscape [19, 20] and leading to the emergence of DTPCs during cancer cell adaptation to therapy has not yet been examined.

The ISR has emerged as a key regulator of resistance to targeted therapy in several cancers [21]. In melanoma, the ISR tunes down the expression of the key transcription factor MITF and thereby promotes de-differentiation and a progressive transition to a mesenchymal-like/invasive cell state [22]. This phenotype switch is accompanied by a decrease in cell proliferation, a metabolic rewiring from glycolysis to OXPHOS and increased resistance to MAPK inhibitors (MAPKi) [23].

Recent studies have linked translational rewiring to the induction of PD-L1 in liver cancer [24], in melanoma [25] and in human lung cancer cells [26]. Additionally, the increase in IDO1 (idolamine 2,3 -dioxigenase) induced by the ISR converts tryptophan to kynurenine, a metabolite with known immunosuppressive function [27]. These recent data indicate that the activation of the ISR pathway may indeed promote immune escape. However, it has also been reported that activation of this pathway by the IFN-*γ* may instead increase cancer cell immunogenicity. Notably, interferon signalling is required for the expression of MHC I [27] and increases the repertoire of putative neoantigens by inducing ribosome stalling and frameshifting [28]. Together these apparently conflicting studies have highlighted the need to better understand the role of ISR in anti-tumour immunity and how to manipulate this pathway to overcome therapy resistance.

Among the strategies used to escape immune surveillance, the assembly of a thick glycocalyx around the cell membrane has been studied in non-cancer systems [29] and linked to infertility [30].

Here we demonstrated that the immune suppressive and immune stimulatory functions of the ISR can be uncoupled by inhibiting a single cancer-specific lncRNA. Retention of this lncRNA at ribosomes following the induction of cap-independent translation, promotes an aberrant translational program promoting the synthesis of PD-L1 and of the glycocalyx and leading to immune suppression. Pharmacological inhibition of this lncRNA, that we named *LISRR*, increases melanoma cell immunogenicity and sensitivity to Immune Checkpoint Blockade (ICB).

## Results

### *LISRR* is associated with polysomes upon induction of the ISR

To gain insights into the common principles governing translational regulation in therapy resistant melanoma, we performed ribosome profiling and RNA sequencing of 3 Patient-Derived Xenograft (PDX) melanoma models, all derived from patients who developed resistance to MAPKi and/or ICB and shown to have engaged the ISR pathway [23]. By comparing the mRNAs and lncRNAs contents in the different ribosomal fractions (Figure 1A-B), we observed that protein-coding RNAs were equally distributed between the ribosomal subunit and polysome fractions (Figure 1A). In contrast, and consistent with their low coding potential, lncRNAs were strongly enriched in the small ribosomal subunit (40S) fraction (Figure 1B). The ISR involves a blockade of translation at the initiation step. Given that the small ribosomal subunit is responsible for the assembly of the initiation complex, we hypothesized that the association of the lncRNA with the 40S subunit may indicate a role in translation initiation.

**Figure 1A-K:**
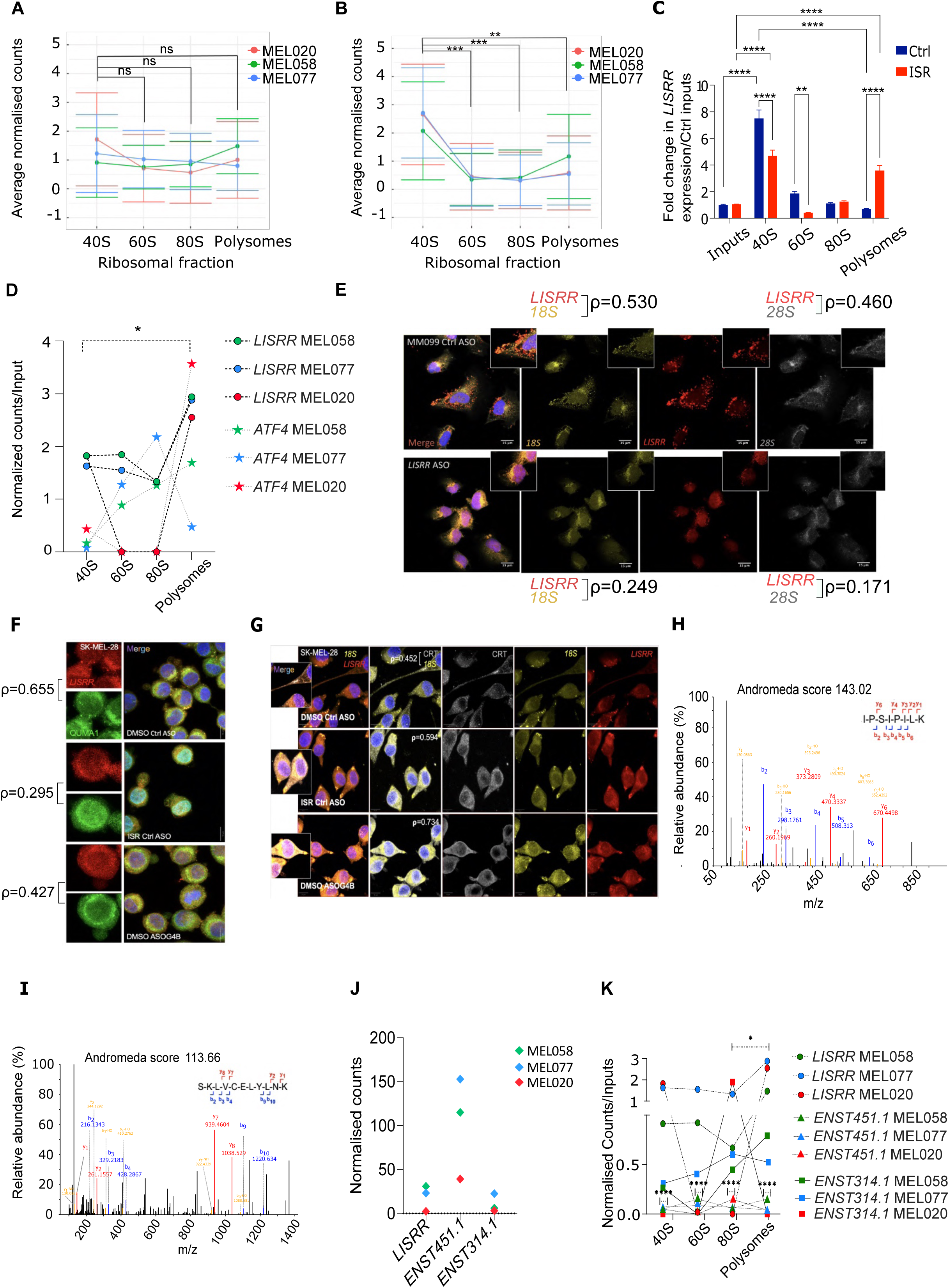
*LISRR* is a lncRNA associated with polysomes during the ISR. **A**. Average mRNA counts in each ribosomal fraction normalized to the average counts of the combined fractions per melanoma model. Significance was calculated by Tukey’s multiple comparisons test. **B**. Average lncRNA counts in each ribosomal fraction normalized on the average counts of the combined fractions per melanoma model. Significance was calculated by Tukey’s multiple comparisons test. **C.** RT-qPCR for *LISRR* expression in different ribosomal fractions. Significance was calculated by two-way ANOVA. **D.** *LISRR* and ATF4 counts per ribosomal fraction normalized on counts in whole lysate (input) in different melanoma PDX models. Significance was calculated with two-way ANOVA. **E.** FISH for *LISRR* (red) and 18S (yellow), 28S (grey) in MM099 in control conditions (Ctrl ASO) and upon *LISRR* KD (*LISRR* ASO). Pearson’s correlation coefficient between the different signals is indicated **F.** FISH for *LISRR* (red) and staining for RNA G4 (QUMA1; green) in SK-MEL-28 in control conditions (DMSO Ctrl ASO), upon induction of ISR (ISR Ctrl ASO) and upon transfection of an ASO blocker recognising *LISRR* G4s (DMSO ASOG4B) **G.** FISH for *LISRR* (red) and 18S (yellow) and calreticulin (grey) of cells in F. **H.** Higher energy collisional dissociation (HCD) spectrum and full annotation of the doubly charged parent ion with m/z = 440.7983 (ENST00000615314.1 - IPSIPILK) in SK-MEL-28 before treatment with salubrinal. Andromeda score is reported as calculated by MaxQuant. Matched fragments are indicated in colour (blue for b-ions, red for y-ions and yellow either for ammonia or water losses). **I.** Higher energy collisional dissociation (HCD) spectrum and full annotation of the doubly charged parent ion with m/z = 683.8756 (ENST00000670451.1 - SKLVCELYLNK) in SK-MEL-28 before treatment with salubrinal. Andromeda score is reported as calculated by MaxQuant. Matched fragments are indicated in colour (blue for b ions, red for y ions and yellow either for ammonia or water losses). **J.** Expression (counts in each fraction normalised on total counts in the whole lysate) of *LISRR*, ENST00000670451.1 and ENST00000615314.1 in different ribosomal fractions derived from PDX models in figure 1A and B. Significance was calculated with two-way ANOVA. **K** Expression (normalised counts) of *LISRR*, ENST00000670451.1 and ENST00000615314.1 in whole lysate from PDX samples depicted in Figure 1A-B. Significance was calculated with two-way ANOVA.

To identify transcripts that are selectively recruited to ribosomes during the establishment of drug-tolerance, we treated SK-MEL-28 melanoma cells with salubrinal, a compound that induces the ISR by inhibiting EIF2⍰ dephosphorylation [22, 24, 25], and performed polysome profiling and RNA-sequencing [23]. Among the candidates significantly associated with the 40S, we identified ENST00000648050.1 and ENST00000650193.1, two transcripts annotated as LINC00941. Although these transcripts do share part of the transcript structure with LINC00941 (ENSG00000235884), they derive from a different gene (ENSG00000285517) (Supplementary Figure 1A). We therefore rename the products of ENSG00000285517 as *LISRR* (for LncRNA ISR Regulated/Regulator). While mainly associated with the 40S in control samples, *LISRR* was strongly enriched at polysomes following ISR activation (Figure 1C). Likewise, *LISRR* was also significantly enriched at polysomes in tumours from the above-described therapy-resistant PDX melanoma models in Figure 1A-B, together with ATF4 (Figure 1D). Localisation of *LISRR* at the ribosomes was further confirmed by single molecule FISH (smFISH; Figure 1E) followed by analysis of expression and co-localisation with rRNAs (Supplementary Figure 1C-D). Accordingly, Pearson’s coefficient (ρ), indicates a slightly higher co-localisation with the 18S that is lost upon knock down of *LISRR* (Supplementary Figure 1C). Analysis of the co-localisation and quantification of the signal further suggests that 60-80% of the ribosomes are occupied by *LISRR* in MM099 at steady state (Figure1G). In silico analysis of *LISRR* sequence with DNA analyser [31] suggests that *LISRR* contains several G-quadruplexes (G4) in its first 400 nucleotides (Supplementary Figure 1E). We therefore designed an ASO blocker (ASOG4B) that could recognise the G4 with highest G4Hunter score. We noticed that the treatment of SK-MEL-28 with ASOG4B, similarly to the ISR activation, induced *LISRR* G4 unfolding (Figure 1F), as indicated by the loss of co-localisation with QUMA1 (Figure 1F and Supplementary Figure 1F), a dye that binds to folded RNA G4. Similarly, in MM099 undergoing ISR, we could not detect any co-localisation between QUMA1 (almost entirely nuclear) and *LISRR* (Supplementary Figure 1E). Interestingly, inhibition of *LISRR* G4 folding in unstressed SK-MEL-28 cells, similarly to ISR induction, triggers endoplasmic reticulum (ER) stress as demonstrated by the strong induction of calreticulin (Figure 1G). Additionally, both the ISR and the ASOG4B increased the ribosome localisation at the ER (Figure 1G and Supplementary Figure 1G).

*LISRR* shift from 40S to polysomes upon induction of the ISR, was unexpected given the low coding potential of *LISRR* predicted by the Coding Potential Calculator (CPC2) algorithm (Supplementary Figure 1H). To rule out the possibility that the transcript is translated we therefore performed a thorough proteomic analysis of small and large peptides in control and salubrinal-treated melanoma cells, which failed to detect any peptide arising from *LISRR* (Supplementary Table I). Conversely, peptides derived from two additional lncRNAs in control samples, but not in salubrinal-treated cells were detected (Supplementary Table I) (Figure 1H-I). Strikingly, while one of the two translated lncRNAs was expressed at very high levels, the other one displayed levels comparable to *LISRR* (Figure 1J). Furthermore, among the three lncRNAs, *LISRR* showed the highest enrichment at the ribosomes (Figure 1K). These data further suggest that our inability to detect *LISRR*-derived peptides is not due to poor sensitivity of our mass spectrometry but rather to the lack of translation or to the production of unstable peptides that are rapidly degraded. This analysis confirmed that the majority of lncRNAs, including *LISRR*, are not translated/unstable in these melanoma cells and provided evidence that the translation of a small minority of lncRNAs is actually repressed upon ISR activation. Importantly, the enrichment of *LISRR* at the polysomes could not be explained by the overall increase in expression of this transcript, as its levels were largely unchanged in cells treated with salubrinal (see input, Figure 1C). Consistently, depletion of ATF4 in salubrinal-treated cells did not impact *LISRR* expression (Supplementary Figure 1I).

*LISRR*, which is a polyadenylated transcript (Supplementary Figure 1J), primarily localizes to the cytoplasm of melanoma cells, as shown by subcellular fractionation (Supplementary Figure 1K) and confirmed by smFISH (Figure 1E). Importantly, its cellular distribution remained unchanged upon the ISR induction (Supplementary Figure 1K).

### *LISRR* expression correlates with resistance to immunotherapy

Analysis of exon expression in normal tissues from the Genotype Tissue Expression (GTEx) indicated that the two transcripts of interest are undetectable in normal adult tissues (Figure 2A), while 3 other transcripts arising from the same locus and extensively studied [32] showed a very weak expression. In the Pan Cancer Atlas (PanCanAtlas) [33] high *LISRR* expression (Figure 2B) and DNA copy number (Figure 2C) is significantly associated with poor patients’ survival. Consistently, *LISRR* expression was significantly higher in metastatic samples compared to primary tumours (Figure 2D).

**Figure 2A-K:**
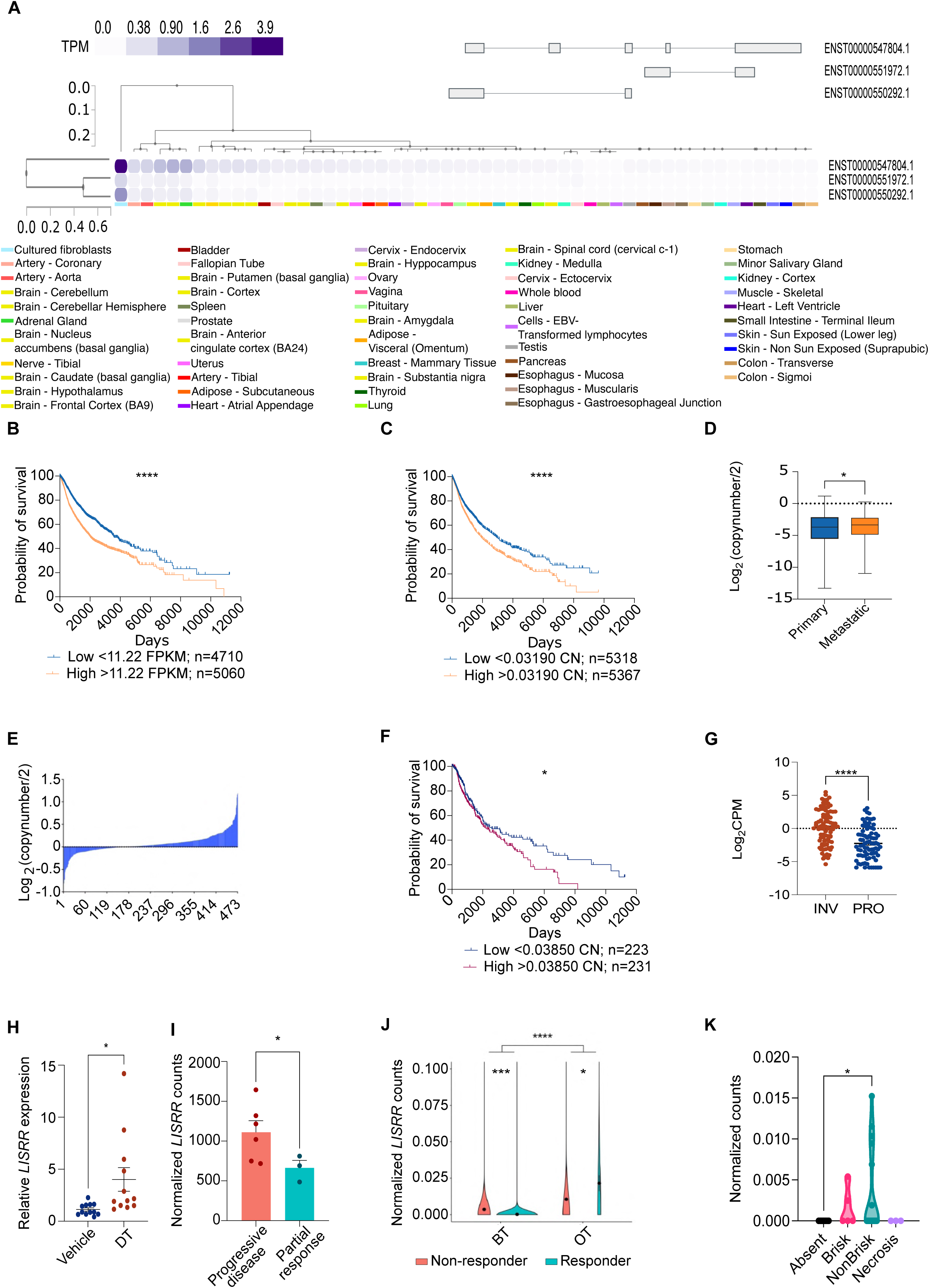
*LISRR* expression and copy number correlate with patient survival and responses to ICB. **A.** Expression of LINC00941 derived transcripts in a panel of adult normal samples from the GTEx database. **B.** Overall survival of patients in the PanCanAtlas cohort as a function of *LISRR* expression Log-rank (Mantel-Cox) test was performed to calculate statistical significance. **C.** Overall survival of patients in the PanCanAtlas cohort as a function of LISRR DNA copy number expressed as Fragments Per Kilobase of transcript per Million reads (FPKM). Log-rank (Mantel-Cox) test was performed to calculate statistical significance. **D.** LISRR DNA copy number in primary and metastatic samples from PanCanAtlas. Unpaired t test was performed to calculate statistical significance. **E.** LISRR DNA copy number amplification in melanoma samples from the TCGA GDC Melanoma dataset. **F.** Overall survival of patients in TCGA GDC Melanoma cohort as a function of LISRR DNA copy number. Log-rank (Mantel-Cox) test was performed to calculate statistical significance. **G.** *LISRR* expression in counts per million (CMP) in TGCA GDC Melanoma dataset with proliferative or invasive signature [30]. Unpaired t test was performed to calculate statistical significance. **H.** Measurement of *LISRR* expression by RT-qPCR in a cohort of melanoma PDX models at relapse after treatment with a vehicle or with targeted therapy (DT). Unpaired t test was performed to calculate statistical significance. **I.** Expression of *LISRR* (read counts normalized by Sleuth) in bulk RNA sequencing libraries derived from BRAF mutant patients partially or not responding to pembrolizumab. Each dot represents aggregate normalized counts of a single patient [36]. Significance was calculated by likelihood ratio test. **J.** *LISRR* expression (read counts normalized by Sleuth) in a melanoma cohort of responders and not responders to immune checkpoint blockade [35], before treatment (BT) and on treatment (OT). Significance was calculated by Mann Whitney test. **K.** Correlation of *LISRR* expression with tumour infiltration status in patients from clinical trial described in J. Significance was calculated by Kruskal-Wallis test.

The *LISRR* locus resides in a region of chromosome 12 (12p11.21) that is amplified in over 60% of melanoma cases (TCGA; Figure 2E). Importantly, the presence of this amplicon was found to be associated with a significantly worse prognosis for melanoma patients (Figure 2F), indicating that one or more genes in this region are important for melanoma progression. Additionally, *LISRR* expression is significantly higher in patients with an invasive signature, associated with resistance to targeted therapy in melanoma TCGA dataset [34] (Figure 2G). *LISRR* expression could be detected in most short-term and established melanoma lines analysed, irrespective of the driver mutation (Supplementary Figure 1J) and in the majority (7 out of 10) of tumours from PDX melanoma models tested (Supplementary Figure 1K), but it is undetectable in Normal Human MElanocytes (NHME; Supplementary Figure 1J). Notably, the highest expression levels were detected in MEL-058 and MEL-083, two models derived from patients progressing on targeted and immune therapy, respectively.

In keeping with this, treating a BRAF-mutant PDX model with a combination of the MAPKi Dabrafenib and Trametinib (DT) promoted emergence of the DTPC population harbouring a mesenchymal-like phenotype [35] and a concomitant increase in *LISRR* expression (Figure 2H). However, no significant correlation between *LISRR* expression and responses to MAPKi could be detected in a large panel of BRAF mutant lines from the CCLE database (Supplementary Figure 2A-B). Instead, a significant negative correlation was found between *LISRR* expression and responses to ⍰-PD-1 in PDX melanoma models (Supplementary Figure 2C-D) and in patients from two independent cohorts [36, 37] (Figure 2I-J). Importantly, in the second cohort, for which single cell RNAseq (scRNAseq) data before (BT) and early on-treatment (OT) are available, *LISRR* expression was found to be induced by the treatment with ICB in cancer cells and to increase in patients refractory to the therapy [36] (Figure 2J). Additionally, in this last cohort, *LISRR* expression is significantly increased in patients with a non-brisk phenotype, characterised by focal immune infiltration and poor responses to ICB (Figure 2K).

These data demonstrate that *LISRR* is a cancer-specific lncRNA whose expression and copy number in melanoma negatively correlate with cancer patients’ overall survival and with response to ICB.

### Inhibition of *LISRR* enhances antimelanoma immune responses in vitro

The negative correlation between *LISRR* expression and response to ICB prompted us to test whether *LISRR* targeting may increase anti-melanoma immune response. We co-cultured IL-2-stimuated HLA matched PBMCs (Figure 3A) with melanoma cells (MM099) harbouring a mesenchymal-like phenotype, exhibiting pre-existing activation of the ISR and high *LISRR* expression [23] (Supplementary Figure 1G). These co-cultures were either exposed to a non-targeting control Antisense Oligonucleotide (Ctrl ASO) or an ASO targeting *LISRR* (Figure 3A-B). As expected, mesenchymal-like melanoma cells untreated or transfected with a non-targeting control were poorly immunogenic [36] (Figure 3A). Interestingly, inhibition of *LISRR* led to an early increase in the PBMC proliferation (Supplementary Figure 3A) and a significant induction of melanoma cell killing (Figure 3C-D), as demonstrated by the decrease in melanoma cell confluency (Figure 3C) and the increase in caspase-3-positive cell counts (Figure 3D). To test whether T-cells are responsible for melanoma cell killing, we activated the PBMCs with a T-cell cocktail, containing ⍰-CD28 and ⍰-CD3, for 48h (Figure 3E-H, Supplementary Figure 3B) before co-culturing them with melanoma cells. Additionally, to rule out any unspecific and/or toxic effect induced by the ASOs, we performed *LISRR* KD with siRNAs (Figure 3F). Again, we could detect an early increase in PBMCs proliferation (Supplementary Figure 3B) as well as increased melanoma cell killing (Figure 3E-H) upon *LISRR* targeting. To rule out that immune cell activation is primed by secretion of soluble factors, we treated PBMCs with conditioned medium derived from M099 melanoma cultures. This did not stimulate PBMC proliferation/activation (Supplementary Figure 3C-D). Accordingly, blocking of MHCI and MHCII with monoclonal antibodies before exposure to HLA-matched PBMCs significantly reduced cell death, indicating that the increase in cancer cell killing upon *LISRR* KD is antigen-dependent (Supplementary Figure 3E).

**Figure 3A-M:**
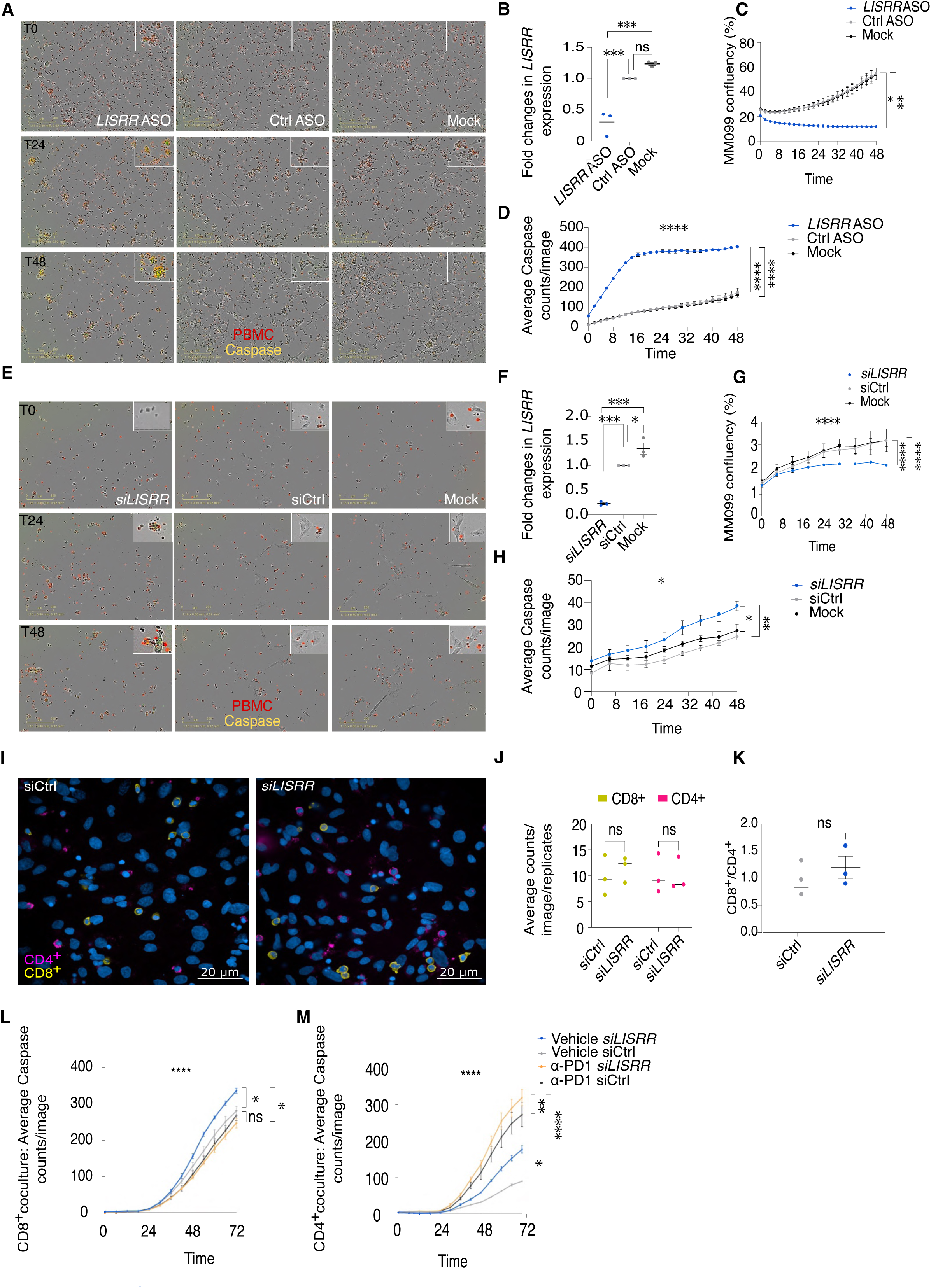
Inhibition of *LISRR* enhances anti-melanoma immune responses. **A.** Representative pictures of HLA-matched PBMCs (red) stimulated with IL-2 and cocultured with MM099 melanoma cell line upon *LISRR* KD using ASOs. Caspase-3^+^ cells are in green/yellow. **B.** Efficiency of *LISRR* KD calculated by RT-qPCR and expressed as fold changes (FC) compared to Ctrl ASO. Statistics calculated by One-way ANOVA. **C.** Analysis of MM099 confluency (%) transfected with *LISRR* ASO, ASO control (Ctrl ASO) or not transfected (MOCK) and co-cultured with PBMCs stimulated with IL-2 only. Statistics calculated Two-way ANOVA mixed effect. **D.** Analysis of Caspase-3 counts in MM099 transfected as indicated in A and cocultured with PBMCs stimulated with IL-2 only. Statistics were calculated by Two-way ANOVA mixed effect. **E.** Representative pictures of MM099 transfected with siRNAs against *LISRR* (*siLISRR*), non-targeting control (siControl) or not transfected (MOCK) and co-cultured with PBMCs (in red) stimulated with a T-cell activating cocktail. Caspase-3^+^ cells are in green/yellow. **F**. Efficiency of *LISRR* KD in experiment in E, calculated by RT-qPCR and expressed as FC compared to Ctrl ASO. **G**. MM099 confluency in the experiment in E. Statistics calculated by Two-way ANOVA mixed effect**. H.** Caspase counts in the experiment in E. Statistics were calculated by Two-way ANOVA mixed effect. **I.** Confocal images of MM099 and PBMCs cocultures in E. In blue nuclei, CD4+ and CD8+ cells are in magenta and in yellow respectively. **J.** Quantification of confocal images from the experiment in I. Average counts of CD4+ and CD8+ per images per biological replicate are shown. Statistics calculated by Two-Way ANOVA. **K.** Ratio CD8+/CD4+ cells from experiment in I. **L.** Caspase counts in coculture of CD8+ cells with MM099 transfected as indicated in E and treated with α-PD1. Statistics calculated by Two-way ANOVA mixed effect analysis. **M.** Caspase counts in coculture of CD4+ cells with MM099 transfected as indicated in E and treated with α-PD1. Statistics calculated by Two-way ANOVA mixed effect analysis.

Cytotoxic T-cells are important effectors in anti-tumour immunity. Although this function has conventionally been ascribed to CD8^+^ cells, intratumoral CD4^+^ T-cells with cytotoxic activity have been recently identified [38].

Analysis of the number of CD8^+^ and CD4^+^ T-cells by immunofluorescence upon *LISRR* inhibition show a trend towards the increase in the amount of CD8^+^ and the CD8^+^/CD4^+^ ratio (Figure 3I-K). Therefore, to further assess the role of CD8^+^ and CD4^+^ T-cells in melanoma cell killing, we isolated CD4^+^ and CD8^+^ T-cells subpopulations from HLA-matched PBMCs and co-cultured them separately with melanoma cells (Supplementary Figure 3F-G). In both cases we could detect Caspase-3 activation (Figure 3L-M) and reduced melanoma cell confluency (Supplementary Figure 3H-I) upon *LISRR* KD, thus suggesting that both MHC I- and MHC II-dependent mechanisms of antigen presentation are at play. However, in CD4^+^ co-cultures the number of caspase-positive cells was approximately half of what was observed in CD8^+^-containing co-cultures unless co-treatment with ⍰-PD1 was applied (Figure 3L-M). All together these results highlight the ability of *LISRR* to protect melanoma cells from T-cell attack.

### *LISRR* regulates translation of selected mRNAs with immunoregulatory functions

To dissect the molecular mechanisms underlying *LISRR*-dependent suppression of melanoma immunogenicity, we silenced the transcript using a pool of siRNAs (Figure 4A) and performed RNA sequencing. While 244 transcripts were significantly (adjusted p<0.05) dysregulated upon *LISRR* KD, only four of them showed a FC>|2|, indicating that *LISRR* KD caused only negligible changes in gene expression. Such fluctuations result from alterations in the stability of specific RNAs, induced by changes in their association with ribosomes, as these transcripts are also differentially translated (Supplementary Figure 4A-B). Gene Set Enrichment Analysis (GSEA; p<0.05 and FDR<0.1), nevertheless, suggested a role for *LISRR* in the regulation of ribosome biogenesis and translation (Figure 4B). We therefore performed polysome profiling and RNA sequencing in *LISRR* KD cells. We found about one fifth of the entire MM099 translatome was affected upon *LISRR* silencing (Figure 4C). As LINC00941 is a divergent transcript of CAPRIN2, a protein involved in RNA transport and translation, we checked whether CAPRIN2 was regulated by *LISRR* KD in our RNA sequencing data. However, this was not the case (these data, provided to the reviewers, have not been included as *LISRR* is encoded by a different gene). Instead, GSEA indicated a significant (nominal p-value=0,022) inverse correlation between the translatome regulated by *LISRR* and mRNAs translated during activation of the ISR (Figure 4D). Particularly relevant was a decrease of ATF4 mRNA at polysomes (Supplementary Figure 4C), which resulted in a significant decrease of ATF4 protein levels as shown by western blot analysis (Figure 4E-F). Similar results were obtained with RNase H-mediated degradation of *LISRR* (Supplementary Figure 4 D-F) and in another mesenchymal-like melanoma line (Supplementary Figure 4G-I).

**Figure 4A-N:**
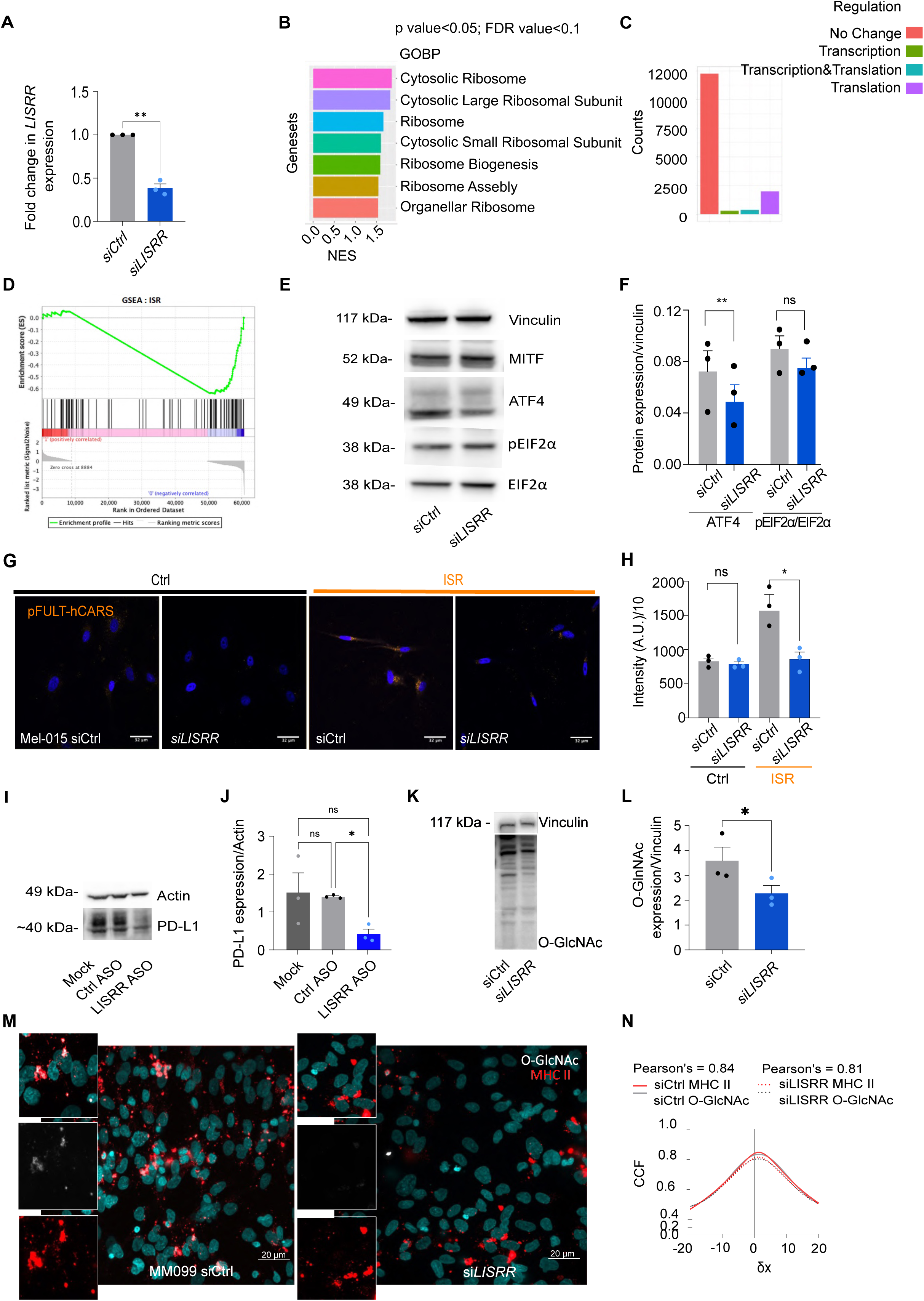
*LISRR* regulates translation. **A.** Efficiency of *LISRR* KD as calculated by RT-qPCR. Values are expressed as fold changes. Paired t test was performed to calculate statistical significance. **B.** GSEA on bulk RNA sequencing data from MM099 cells upon *LISRR* KD. Threshold values p-value<0.05; FDR-val<0.1. **C.** Effect of *LISRR* KD on the transcriptome and translatome as calculated by Ribosomal Investigation and Visualization to Evaluate Translation (RIVET) [82] analysis on bulk RNA sequencing data from polysome profiling of MM099 cells upon *LISRR* KD. **D.** Correlation between bulk RNA sequencing data from ribosome profiling of MM099 cells upon *LISRR* KD and the ISR translational signature by GSEA (nominal p-value = 0.02259887). **E.** Western blot for MITF, ATF4, peIF2α/eIF2α upon *LISRR* KD. Vinculin was used as loading control. **F.** Densitometric analysis of the western blot in F. Paired t test was performed. **G.** Confocal images showing Mel-015 cells infected with pFULThCARS reporter in control condition (DMSO) and upon ISR induction (salubrinal). Nuclei are in blue and pFULThCARS is in orange. **H.** Quantification of luciferase activity of pFULT-hCARS reporter in Mel-015 cells upon *LISRR* KD and ISR activation. **I.** Representative western blot for PD-L1 in MM099 upon *LISRR* KD. Actin was used as a loading control. **J.** Densitometric analysis of the western blot in I. Statistics were calculated by paired t-test. **K.** Representative western blot for O-linked N-acetylglucosamine in MM099 upon *LISRR* KD. Vinculin was used as a loading control. **L.** Densitometric analysis of western blot in K. Statistics were calculated by paired t-test. **M.** Immunostaining for MHCII and O-linked N-acetylglucosamine (grey) in melanoma cell (red). **N.** Evaluation of co-localisation between MHCII and O-linked N-acetylglucosamine as measured by Cross Correlation Function (CCF) with a pixel shift of δ = ±20.

To further substantiate these findings, we built an ATF4-reporter system (pFULT-hCARS, Supplementary Figure 4J), in which the dt-Tomato and luciferase genes were placed under control of the human ATF4-responsive CARS promoter [39]. The reporter system was introduced into a PDX-derived melanoma cell line (Mel-015). As expected salubrinal exposure led to a robust activation of the reporter in cells transfected with non-targeting siRNAs (siCtrl). In contrast, co-transfection with *LISRR*-targeting siRNAs (*siLISRR*) led to a significant decrease in luciferase and fluorescent reporter activity (Figure 4G-H and Supplementary Figure 4K).

The further mining of our polysome profiling sequencing dataset highlighted two additional important findings.

First, translation of *PD-L1* was found to be dependent on *LISRR*, as its KD decreased *PD-L1* mRNA expression and early polysome occupancy in ISR-activated cells (Supplementary Figure 4L). Downregulation of PD-L1 at the protein level could also be validated by western blotting (Figure 4I-J).

Additionally, GSEA analysis revealed a translational downregulation (p-value<0.05; FDR-val<0.1) of genes involved in the biosynthesis and metabolism of glycolipids and glycoproteins upon *LISRR* KD (Supplementary Figure 4M). This was further confirmed by the decrease in O-GlcNac detected by western blotting (Figure 4K-L) in MM099 transfected with siRNAs (Supplementary Figure N) against *LISRR* or with a *LISRR*-targeting ASO (Supplementary Figure 4O-P). Similar results were obtained in MM165, another mesenchymal-like melanoma line (Supplementary Figure 4Q-R). Additionally, co-staining for the MHCII and O-GlcNac revealed an overlap between the glycosylation signal and the antigen presenting machinery (Figure 4M-N). Given the previously reported negative association between the built up of the glycocalyx and immunogenicity of tumours [40], *LISRR* may promote immune evasion by inducing the presentation of sugars at the surface of cancer cells.

These data suggest that *LISRR* retention at polysomes in cells undergoing ISR activation facilitates the translation of selective transcripts, such as *PD-L1* and genes involved in biosynthesis and metabolism of glycans. As such, they provide mechanistic insights into how *LISRR* regulates melanoma immunogenicity.

### Recruitment of DAZAP-1 to ribosomes elicit *LISRR*-driven ISR activation

To understand how *LISRR* recruits specific transcripts to the ribosome during adaptation induced by the ISR, we compared the RNA binding protein footprint in transcripts enriched and depleted at ribosomes upon *LISRR* KD with oRNAment [41] This search indicated an enrichment in binding motifs (Supplementary Figure 4S) for a protein called DAZAP1 (Deleted in Azoospermia Associated Protein 1) in the transcripts depleted from polysomes upon *LISRR* KD (Figure 5A). Most of the transcripts arising from LISRR locus contain such binding sites (Supplementary Figure 5A). DAZAP1 partners (DAZ proteins) are hosted on the Y chromosome in a region deleted in infertility. By mining publicly available scRNAseq data of patients with Non-Obstructive Azoospermia (NOA; GEO accession number GSE153947), we found that low levels of DAZAP1 correlate with loss of PD-L1 expression (Supplemental Figure 5B-C) and glycosylation defects (Supplemental Figure 5D).

**Figure 5A-O:**
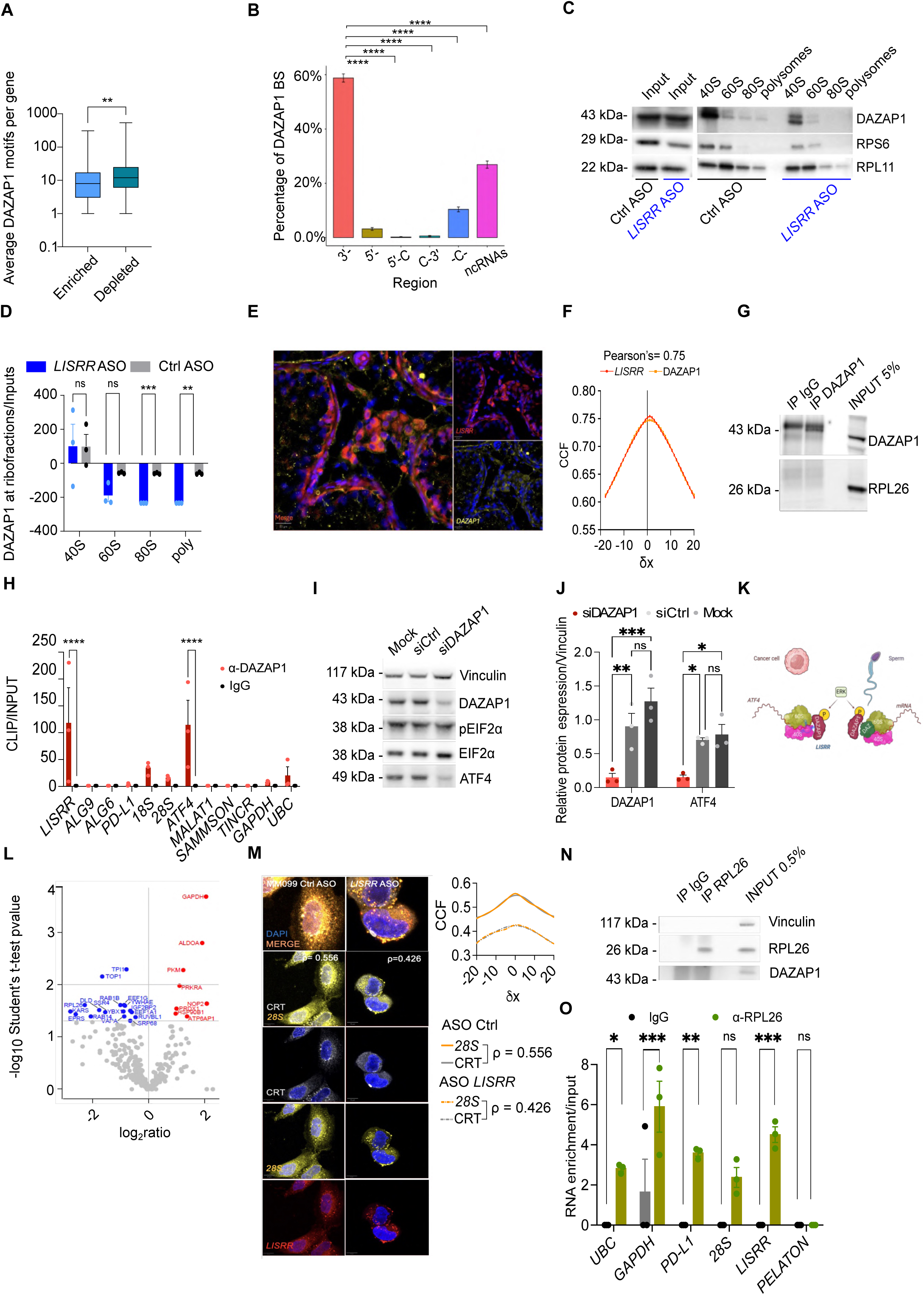
*LISRR* affects ribosome composition and interactions. **A.** Average number of DAZAP1 binding motifs per transcript in mRNA significantly enriched or depleted at polysomes upon *LISRR* KD. **B.** Distribution of DAZAP1 binding sites in mRNAs dysregulated by *LISRR* KD. Significance was calculated by Wilcoxon test. **C.** Representative western blot for DAZAP1 in MM099 ribosome profiling experiment upon *LISRR* KD. Ribosomal proteins are used as a loading control. **D.** Densitometric analysis of the western blot in C. **E.** Staining for DAZAP1 (yellow) and *LISRR* (red) in testicular sperm extraction samples from UZ Leuven. **F.** Co-localisation between DAZAP1 and *LISRR* as measured by Cross Correlation Function (CCF) with a pixel shift of δ = ±20 **G.** Efficiency of DAZAP1 CLIP as assessed by western blot for DAZAP1 and for Ribosomal proteins in MM099. **H.** RT-qPCR for *LISRR* and *LISRR* translational targets in MM099 DAZAP1 CLIP. **I.** Western blot for ATF4 and ISR modulators in MM099 upon *LISRR* KD. **J.** Densitometric analysis of the blot in I. Statistics were calculated by ANOVA. **K.** Model of *LISRR*-DAZAP1 interaction at the ribosomes. **L.** Volcano Plot indicating by log2 ratio the proteins enriched and depleted at ribosomes upon *LISRR* KD**. M.** FISH for *LISRR* (red) and 28S (yellow), calreticulin (grey) in MM099 in control conditions (Ctrl ASO) and upon *LISRR* KD (*LISRR* ASO). Pearson’s correlation coefficient between the different signals is indicated **N.** Efficiency of RPL26 RIP as assessed by western blot for RPL26, DAZAP1 and Vinculin in MM099. **O.** RT-qPCR for *LISRR* in MM099 RPL26 RIP. 28S and other translated mRNA are used as positive controls The unrelated cytoplasmic lncRNA PELATON is used as a negative control.

As loss of the Y chromosome (LoY) was recently associated with immune evasion and exhaustion [42], we thought to check whether correlation between survival and *LISRR* expression was dependent on the presence of the Y chromosome. Accordingly, correlation was abolished upon loss of Y chromosome (Supplemental Figure 5E-F). These data suggest that in the absence of DAZAP1, *LISRR* may become dispensable for the progression of those tumours. DAZAP1 is a protein phosphorylated by ERK and recruited at ribosomes through germ cell-specific DAZ proteins with a role in cap-independent translation [43]. DAZAP1 contains 2 disordered and 2 RRM domains and thus it could bind *LISRR* and mRNA at the same time. We therefore analysed its association with ribosomes upon *LISRR* KD. Consistently, polysome profiling followed by western blot indicated that *LISRR* downregulation results in a significant dissociation of DAZAP1 from monosomes and polysomes (Figure 5C-D). Furthermore, expression of *LISRR* and its co-localisation with DAZAP1 could be detected in the cytoplasm of stromal cells in the testis of patients with NOA (Figure 5E-F). However, we could detect only a weak, although significant, correlation between translation rates and presence of DAZAP1 BS in the transcripts depleted from polysomes upon *LISRR* KD, indicating that this RNA binding protein may not explain entirely the translational changes exerted by *LISRR* (Supplementary Figure 5G). In keeping with this, UltraViolet crosslinking and immunoprecipitation (CLIP) with a DAZAP1 specific antibody (Figure 5G) followed by RT-qPCR confirmed that DAZAP1 binds to *LISRR* and to *ATF4* mRNA, but not *PD-L1*(Figure 5H). In keeping with this, DAZAP1 KD with a pool of siRNAs, resulted in a significant ATF4 downregulation thus phenocopying the effect of *LISRR* KD on ATF4 (Figure 5I-J). These data indicate that, upon *LISRR* recruitment, DAZAP1 regulates the translation of *ATF4* which in turn triggers the ISR (Figure 5K). The effect of DAZAP1 on translation are therefore secondary to the induction of *ATF4* translation.

### Inhibition of *LISRR* alters the assembly of the ribosomes

Since PD-L1 and some of the glycosylases found to be regulated by *LISRR* were not binding directly to DAZAP1 nor were they regulated directly by the ISR and thus not CAP-independent (Supplementary Figure 5H-J), we tested whether *LISRR* can directly affect the composition of the ribosomes. Towards this, we isolated the ribosomes on a sucrose gradient and performed mass spectrometry in control samples and upon *LISRR* KD (Table II and Figure 5L). Downregulation of *LISRR* resulted in the dissociation from the ribosomes of the ribosomal protein RPL26, abundantly expressed in the testis [44]. RPL26 is located near the ribosome exit tunnel [45] where it plays a role in co-translational translocation of proteins to the ER. In keeping with this, several proteins belonging to the TRAM/TRAP complex on the cytosolic side of the endoplasmic reticulum (ER) were also depleted from the ribosomes (Figure 5L). Importantly, among them we detected SSR4, which is mutated in a congenital disorder of glycosylation [46]. Consistently, inhibition of *LISRR* in MM099 reduces the anchoring of the ribosomes to the ER as demonstrated by the decreased co-localisation between *LISRR*, 28S and the calreticulin (Figure 5M). Sensitivity to treatment with 1,6 hexanediol suggested that this interaction, taking place upon G4 unfolding during the ISR, involves phase separation (Supplementary Figure 6A). Lastly, RIP for RPL26 (Figure 5N) retrieved specifically *PD-L1* and *LISRR* indicating that the recruitment of the large ribosomal subunit occurs through this protein (Figure 5O). These results indicate that the defective glycocalyx and PD-L1 expression [47] are likely to be caused by a profound defect in protein trafficking and glycosylation due to a defective anchoring of the ribosomes to the ER upon *LISRR* depletion.

### Systemic inhibition of *LISRR* affects immune responses in vivo and re-sensitizes to ⍰-PD1 therapy

To assess the clinical relevance of these findings in vivo, we engrafted a PDX melanoma model resistant to immunotherapy into immunocompromised mice (NOG) as well as into mice humanised with CD34^+^ hematopoietic stem cells (hu-NOG; Supplementary Figure 6B). Systemic inhibition of *LISRR* by subcutaneous injection of an ASO (Figure 6A) in tumour-bearing immune compromised mice did not affect overall survival (Figure 6B), indicating that *LISRR* KD is not required per se for the in vivo growth of melanoma cells. In humanised mice, however, the same treatment significantly delayed tumour growth (Figure 6C) and resulted in increased overall survival (Figure 6B). Critically, *LISRR* targeting re-sensitized this PDX model to ICB (Figure 6D). Western blot for O-linked N-acetylglucosamine confirmed its downregulation *in vivo* upon pharmacological inhibition of *LISRR* (Supplementary Figure 5C-D).

**Figure 6A-M:**
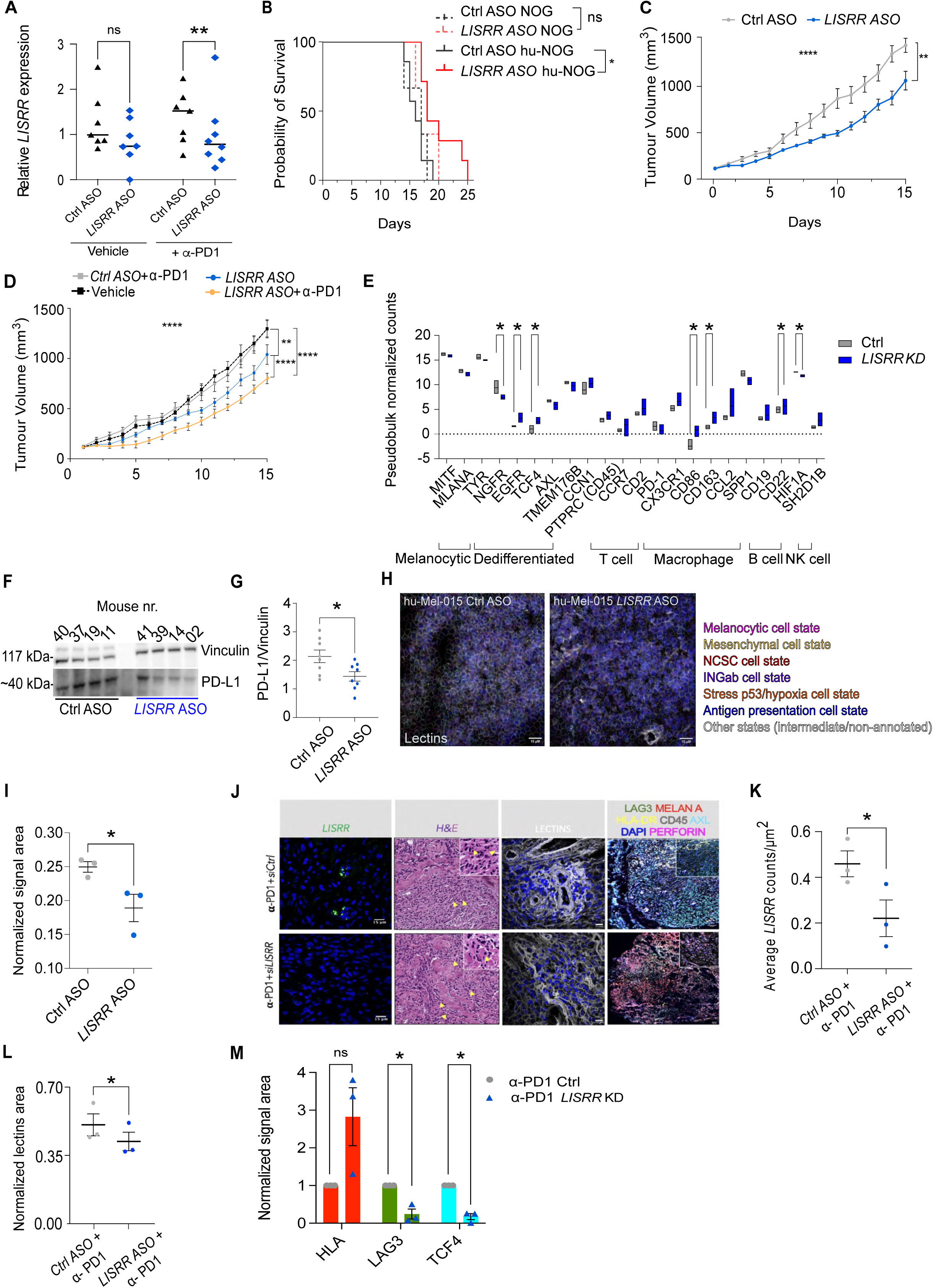
Inhibition of *LISRR* overcomes primary resistance to ICB. **A.** Efficiency of *LISRR* KD in vivo in PDX models upon systemic administration of an ASO against *LISRR* or a non-targeting control. Multiple t test was performed to calculate statistical significance. **B**. Kaplan-Meier plot of immune deficient and immunocompetent mice upon systemic inhibition of *LISRR*. **C.** Comparison of the tumor growth over time of humanized PDX melanoma models treated with an antisense against *LISRR* (*LISRR* ASO n=7), a non-targeting control (Ctrl ASO n=7) or a vehicle (n=5). Two-way ANOVA mixed effect was performed to calculate statistical significance. **D.** Comparison of the tumor growth over time of humanized PDX melanoma models treated with an antisense against *LISRR* (*LISRR* ASO n=7), with α-PD1 (n=6), or with a combination of the two (ASO+α-PD1 n=5). Two-way ANOVA mixed effect was performed to calculate statistical significance. **E.** Quantification of immune- and melanoma drug-tolerant-markers upon *LISRR* KD, as assessed by smFISH and molecular cartography in hu-PDX tumour sections described in C. **F.** Western blot quantifying the expression of PD-L1 in PDX tumours depicted in D. **G.** Densitometric analysis of the western blot in F. Statistical significance was calculated by unpaired t test. **H.** Staining for lectins in hu-PDX tumour sections described in C and its quantification. The segmentation indicates the melanoma state of the cells. Nuclei are labelled with DAPI in blue. Lectins are in grey. **I.** Quantification of the picture in H. Statistical significance was calculated by unpaired t-test. **J.** Image of the KD of *LISRR* in combination with α-PD1 in PDTF derived from an NRAS mutant patient progressing on anti-PD1. Left panel: smFISH for *LISRR*. Central panel: H&E staining of the slide immediately adjacent to right panel. Yellow arrows indicate the lymphocytes infiltrating the tissue. Right panel: lectins staining. Extreme right: Akoya phenocycler staining for the indicated melanoma and immune markers**. K.** Quantification of *LISRR* KD from the smFISH in the left panel in J. **L.** Quantification of the lectins staining shown in the right panel in J. Statistics were calculated by paired t-test. **M.** Quantification of melanoma and immune markers upon *LISRR* inhibition in 3 different PDTF. Statistics were calculated by paired t-test.

To further study the molecular and cellular consequences of *LISRR* KD in vivo, we profiled the melanoma lesions using a highly sensitive spatial transcriptomics method (Molecular cartography, Resolve Biosciences). This analysis confirmed the successful targeting of *LISRR* (Figure 6E) and revealed that *LISRR* KD profoundly affects melanoma heterogeneity by significantly reducing the amount of drug-tolerant subpopulations, as demonstrated by the decrease in several drug-tolerant markers (Figure 6E) including NGFR. Increased infiltration of immune cells was also detected (Figure 6E), together with a decrease in T-cell exhaustion markers (Figure 6E). Western blot for PD-L1 on these samples confirmed its downregulation upon *LISRR* KD (Figure 6F). Lastly, downregulation of the glycocalyx upon *LISRR* KD was also detected in hu-Mel-015 by lectin staining (Figure 6H). Similar results were obtained by inhibiting *LISRR* expression for 48h with siRNAs and ASO in tumour explants derived from melanoma patients (n=3) progressing on immune checkpoint blockade (Figure 6J). Staining with Akoya phenocycler of the tumours upon co-treatment with anti-PD-1 and *LISRR* KD revealed a marked decrease in dedifferentiated cells (AXL and/or TCF4 positive), a decrease in LAG3 expression and marked increase in HLA-DR and CD45 positive cells (Figure 6J-M). In keeping with this, H&E staining shows a remarkable increase in lymphocytes’ size, compatible with their maturation (Figure 6J). Strikingly, staining of the glycans with lectins also revealed a clear decrease in their exposure on the glycocalyx upon *LISRR* KD in vivo (Figure 6K).

These data provide a proof of concept that inhibition of *LISRR* re-sensitises resistant tumours to ICB.

## Discussion

We describe herein a role for *LISRR* as a critical regulator of melanoma immunogenicity.

We show that this non-coding transcript is widely expressed in cancer/melanoma cells, but not in adult normal cells, to promote an immune-tolerant translatome. While evidence of specific ribosomal proteins contributing to ribosome heterogeneity is emerging [44], our data provide evidence that tailor-made ribosomes can be obtained also by incorporating individual lncRNAs into the ribosome. We had previously demonstrated that transcripts engaged in the ISR are enriched for G4 [23]. Here we show that *LISRR* G4 are unfolded in cells undergoing ISR and that their artificial unfolding in unstressed cells is sufficient to generate ER stress. The retention of *LISRR* at polysomes during the ISR, primed by the unfolding of its G4 and possibly leading to the recruitment of RPL26 through multivalent interactions, generates cancer-specific ribosomes and contributes to the emergence of DTPCs. Mechanistically, this process favours translation of mRNAs with immunosuppressive functions, such as that encoding for the immune checkpoint receptor PD-L1 and several enzymes implicated in the formation of the glycocalyx. This carbohydrate sugar coat acts as a physical barrier to protect cancer cells from immune recognition and attack [48], and its aberrant composition is a hallmark of cancer currently in use as predictive biomarker [49]. The assembly of a thick glycocalyx on the cell membrane as a mean to escape the immune system is a strategy shared by many biological systems. Spermatozoa for instance use it to safely travel through the seminal tubules during capacitation and later on through the female reproductive tracts [29]. As a matter of fact, mutations in genes encoding for essential mediators of the glycocalyx deposition result in infertility [30]. Modified sugar analogues have been designed to target glycosylation, however, given their limited uptake by the cells, these inhibitors are of limited therapeutic benefit. Our data offer an alternative, clinically compatible, approach to target this pathway in a cancer cell-specific manner. Consistently, we show that targeting *LISRR* sensitizes refractory melanoma to immune checkpoint blockade. Given that *LISRR* copy number and expression are increased in many different cancer types, we propose that this approach may be applicable beyond melanoma. From a fundamental point of view, our data support a model in which a single cancer cell-specific lncRNA, namely *LISRR*, can participate to the assembly of specialized cancer ribosomes to promote immune evasion and thereby tumour progression and resistance to immunotherapy. Specifically, to rewire translation *LISRR* hijacks the RNA-binding protein DAZAP1. In normal cells, DAZAP1 is recruited to the ribosome by members of the DAZ family of proteins [43] that are almost entirely germ-cell-specific [49] and definitely not expressed in melanoma. The deletion of the highly conserved members of this family has been associated with infertility [50], however, only an handful of its translational targets are known [51]. Our study sheds new light on the potential role of DAZ proteins in infertility through the identification of ATF4 and the ISR as direct targets. Furthermore, we linked the deficiency of DAZAP1 in patients with NOA to defective glycosylation and impaired immune checkpoint blockade. This highlights the evolutionary convergence between sperm and cancer cells in using the glycocalyx as a mechanism to shield from the immune response.

Lastly, since *LISRR* is primate-specific, this study provides yet another example of non-conserved transcript that contribute to the development of a human disease. Conservation has long been used to imply functionality of (lnc)RNAs [52] and this has led to a bias in the recognition of important (non-)coding drivers in cancer [53]. However, since cancer cells are exposed to a much higher selective pressure compared to normal cells, especially when exposed to therapeutic pressure, it is reasonable to assume that these cells may evolve de novo survival mechanisms that involve portions of the genome that are not necessarily functional in normal cells [54].

### Limitations of the study

This study does not aim to develop a market-ready anti-LISRR drug. Although our conclusions are based on several orthogonal assays, numerous ASOs and/or siRNAs need to be tested to achieve such a drug, with thorough evaluation of toxicity and off-target effects. Furthermore, comparison of data obtained *in vivo* and *ex vivo,* suggest that naked ASO delivery is inefficient and thus not amenable for the treatment of patients and would thus require the development of lipid nanoparticles.

## Material and methods

### Cell lines and transfection

SK-MEL-28 (from ATCC) were grown in RPMI 1640 (Gibco BRL Invitrogen), supplemented with 10% FBS (Gibco BRL Invitrogen) and 1% penicillin/streptomycin (Sigma-Aldrich). The patient-derived MM cell lines (a gift from G.-E. Ghanem) were grown in F-10 (Gibco BRL Invitrogen), supplemented with 10% FBS (Gibco BRL Invitrogen) and 1% penicillin/streptomycin.

All cell lines used are of human origin and were confirmed negative for mycoplasma before experimental use by using the MycoAlert Mycoplasma Detection Kit (Lonza) according to the manufacturer’s specifications.

Gender of the patients from whom the cell cultures were derived is as follows (female: F; male: M): SK-MEL-28, M; MM034, F; and MM011, F; MM099, M; MM165, M; MM001, F; MM074, F; MM087, F; A375, F; MM031, M; MM162, M; MM047, M; MM057, F; MM163, F; MM052, M; MM029, M.

For ATF4 knock-down, melanoma cells were plated in 6-well plates and transfected with Lipofectamine 2000 (Thermo Fisher Scientific) according to manufacturer instructions either with 25 nM non-targeting siRNA control or with targeting siRNA (SMART-pool, Dharmacon) 24h after seeding. For *LISRR* KD either control (MISSION® siRNA Universal Negative Control #1 and #2, Sigma Aldrich) or *siLISRR* (Sigma Aldrich, for sequences refer to Table S1) were used at a concentration of 25 nM. For ASO-mediated KD the ASO against *LISRR* (Table S2) or a non-targeting control (Table S2) were used at a final concentration of 50 nM. Cells were collected for RNA and protein extraction 48h after transfection.

### ASO synthesis

Antisense oligonucleotides synthesis was performed on an Expedite 8909 DNA synthesiser (Applied Biosystems) by using the phosphoramidite method. The oligomers in the form of phosphorothioated LNA GapmeRs and fully modified LNA blocker, were deprotected and cleaved from the solid support by treatment with AMA solution (1:1 mixture of ammonia 33% and methylamine 40%) for 2h at 40°C. After gel filtration on an illustra NAP-25 column (Sephadex G-25 DNA Grade; GE Healthcare) using water as an eluent, the crude mixture was purified using a MonoQ HR 10/100 GL anion exchange column (GE Healthcare) with the following gradient system: 10 mM NaClO_4_ and 20 mM Tris-HCl in 15% CH_3_CN, pH 7.4 (A); 600 mM NaClO_4_ and 20 mM Tris-HCl in 15% CH_3_CN, pH 7.4 (B), 0-80% buffer B in 40 minutes, 2 mL/min. The low-pressure liquid chromatography system consisted of a Primaide PM1110 HPLC pump, Mono-Q HR 10/100 GL column, and Primaide PM1410 HPLC UV detector. The product-containing fractions were desalted on a NAP-25 column and lyophilized, and ASO sequences were analysed by mass spectrometry.

### Coculture experiment

Upon knock-down of *LISRR*, MM099 were cocultured with HLA matched PBMCs (Precision for medicine) at 1:5 ratio. MM099 were seeded in a 96-well plate and transfected 24h after using lipofectamine LTX (Thermo Fisher Scientific), according to manufacturer instruction. Pre-activated PBMCs were added 48h post-transfection and the cocultures were followed for up to 120h.

To activate the T-cells, freshly thawed PBMCs were cultured in RPMI 1640 (Gibco BRL Invitrogen), supplemented with 10% FBS (Gibco BRL Invitrogen) and 1% penicillin/streptomycin (Sigma-Aldrich) and 3 µg/mL of human ⍰-CD3 (Thermo Fisher Scientific), 5 µg/mL of human ⍰-CD28 (Thermo Fisher Scientific), and 200 ng of IL-2 (Immunotools) for 48h before coculturing them with melanoma cells.

Cocultures were monitored by use of IncuCyte ZOOM system (Essen BioScience), four images per well were taken every 2h. PBMCs were labelled with 1:1000 Incucyte® Nuclight Rapid Red Dye for Live-Cell Nuclear Labeling Incucyte (Sartorius) while cells death was measured by adding the IncuCyte Caspase 3/7 Green Apoptosis Assay Reagent 1:5000 to the cocultures (Essen BioScience). The cell confluency and caspase positivity were measured and analysed with IncuCyte ZOOM software. The same experiment was performed with the same settings using ASOs.

In order to block MHC I or MHC II, MM099 were cultured with ⍰-human MHC class I HLA-A, HLA-B, HLA-C (#78792101, clone W6/32; inVivoMab™) or ⍰-human MHC class II HLA-DP/DQ/DR (#47AU4, clone IVA12; antibodies online) at a final concentration of 10 µg/mL before seeding. After 48h, the antibodies were refreshed by supplementing the media with an additional 10 µg/mL of either one or the other.

For DAZAP1 KD, MM099 were seeded in 6-well plates and transfected with 50 nM of sipool DAZAP1 or siCtrl (Dharmacon) using LIPO LTX according to manufacturer instructions. The day after the first transfection a second transfection was performed at the same concentration. Cells were collected 72h after the first transfection and processed for RNA extraction and qPCR and protein extraction and Western Blot.

### Polysome profiling

Melanoma cells for polysome profiling were plated in order to obtain a confluency of 70% on the day of the experiment. MM099 (plated in 15 cm dishes, 2 per condition) were transfected using lipofectamine 2000 (Thermo Fisher Scientific), according to manufacturer instruction, either with 25 nM sipool control (MISSION® siRNA Universal Negative Control #1 and #2, Sigma Aldrich) or with 25 nM of sipool for *LISRR* (Sigma Aldrich, for sequences refer to table) for 48h. At endpoints, cells were treated with 100 μg/mL of cycloheximide (Sigma-Aldrich) for 12 minutes at 37 °C, collected, and resuspended in lysis buffer 2 [20 mM Trish-HCl pH 7.4, 10 mM NaCl, 3 mM MgCl2 supplemented with 1.2% TRITONX-100, 0.2 M Sucrose, 100 μg/mL cycloheximide, 20 U/μl SUPERase-IN RNase Inhibitor (Invitrogen, Thermo Fisher Scientific), supplemented with fresh 1× Halt Protease and Phosphatase Inhibitor Single-Use Cocktail (Life Technologies)]. Lysates were then incubated agitating at 4°C for 35 minutes, and then centrifuged at 17,000 RCF for 15 min at 4°C. Lysates were loaded on a sucrose gradient (15%-50%) prepared in buffer G (20 mM Tris-HCl, 100 mM KCl, 10 mM MgCl2 supplemented with 1 mM DTT (Sigma Aldrich) supplemented with fresh 100 μg/mL cycloheximide. Samples were then centrifugated in an SW41Ti rotor (Beckman Coulter) at 37,000 RPM at 4 °C for 150 minutes. The fractions were obtained with a Biological LP System (Bio-Rad). A total of 14 (600 μl) fractions were collected per sample and pulled together to obtain clean 40S, 60S, 80S, and polysomes.

Polysome profiling of PDX-derived samples, was performed as previously described [23] with minor adjustments. Samples homogenized in lysis buffer were loaded onto a 5%-20% linear gradient and centrifugated at 37,000 RPM for 170 minutes at 4 °C in an SW41Ti rotor (Beckman Coulter).

### UV Crosslinking and RNA ImmunoPrecipitation

MM099 were plated in 15 cm dishes. At endpoint, crosslinked by irradiating them with 4 mJ/cm^2^ UV (265nm) using a UVP crosslinker (Analytik Jena, France). After lysis in immunoprecipitation buffer [20 mMTris-HCl pH 8.0, 200 mM, MgCl2 2.5 mM. TRITONX-100 1%, 1mM DTT (Sigma), 20 U/μl SUPERase-IN RNase Inhibitor (Invitrogen, ThermoFisher Scientific), supplemented with fresh 1× Halt Protease and Phosphatase Inhibitor Single-Use Cocktail (Life Technologies), the cells were then incubated for 1h at 4°C on a rotating wheel and then spun down at 17000 RCF for 15 minutes at 4 °C. Supernatant was collected and protein content measured with Bradford assay (Bio-Rad). 5% of the total lysate was kept respectively as RNA and protein input. A minimum of 1 mg of protein was used for the RIP: DAZAP1 was immunoprecipitated using 6 µg of specific antibody (α-DAZAP1, A303-985A-T, Bethyl Laboratories) overnight at 4 °C on a rotating wheel. 6 µg of normal rabbit IgG (12-370, LOT: 3493998, Millipore) was used as control. The lysate was then coupled to 60μl of Protein G Dynabeads (Invitrogen) and left 4h at 4 °C on a rotating wheel. Beads where then washed several times with immunoprecipitation buffer. For the crosslinked samples, beads were washed also several times with DEPC water (Invitrogen). RNA and proteins were collected afterwards. The experiment was validated through qPCR and western blot. For RPL26 RIP 2mg of MM099 extracts were incubated for 3 h and 30’ with 6 µg of α-RPL26 (A300-686A sanbio) or normal rabbit IgG conjugated to 80 µl of Protein A Dynabeads (Invitrogen).

The relative expression of the genes of interest for the CLIP was calculated applying the following ΔΔCt method: the Ct value of the CLIP was subtracted from the Ct value of the Input for every gene, thus obtaining the ΔCt for each gene in the CLIP sample. The CLIP ΔCt was then subtracted from the ΔCt of the immunoglobin (IgG) for every gene, thus obtaining the ΔΔCt. To calculate the fold enrichment, the following equation was applied: fold enrichment= −ΔΔ 2 Ct.

### MS-based proteomics analysis

Both control and salubrinal-treated cell pellets were digested using the PreOmics iST sample preparation kit following the manufacturer’s guidelines. In all cases, proteolytic peptides were separated by reversed-phase chromatography on an EASY-nLC 1200 ultra-high-performance liquid chromatography (UHPLC) system through an EASY-Spray column (Thermo Fisher Scientific), 25 cm long (inner diameter 75 µm, PepMap C18, 2 µm particles), which was connected online to a Q Exactive HF (Thermo Fisher Scientific) instrument through an EASY-Spray™ Ion Source (Thermo Fisher Scientific). Both for library and study samples, the purified peptides were loaded in buffer A (0.1% formic acid in water) at constant pressure of 980 Bar. They were separated through the following gradient: 30 min of 3-23% of buffer B (0.1% formic acid, 80% acetonitrile), 5 min 23-30% of buffer B, 1 min 60-95% buffer B, at a constant flow rate of 250 nl/min. The column temperature was kept at 45°C under EASY-Spray oven control. The mass spectrometer was operated data-dependent acquisition (DDA) mode. Briefly, MS spectra were collected in the Orbitrap mass analyzer at a 60,000 resolution (200 m/z) within a range of 300–1550 m/z with an automatic gain control (AGC) target of 3e6 and a maximum ion injection time of 20 ms. The 15 most intense ions from the full scan were sequentially fragmented with an isolation width of 1.4 m/z, following higher-energy collisional dissociation (HCD) with a normalized collision energy (NCE) of 28%. The resolution used for MS/MS spectra collection in the Orbitrap was 15,000 at 200 m/z with an AGC target of 1e5 and a maximum ion injection time of 80 ms. Precursor dynamic exclusion was enabled with a duration value of 20s. MS Raw files were processed with MaxQuant (MQ) version 2.0.3.0 integrated with Andromeda search engine. Sequenced non-coding RNAs were in silico translated into amino acid sequences already in FASTA format, using the “Six-frame translation” tool available in the MQ suite. In parallel, MSMS spectra were also searched against the human reference proteomes (Uniprot UP000005640, 80,027 entries). The search included cysteine carbamidomethylation as a fixed modification and methionine oxidation and acetylation of the protein N-terminus as variable modifications. Required minimum peptide length was 7 amino acids and maximum mass tolerances were 4.5 p.p.m. for precursor ions after nonlinear recalibration and 20 p.p.m. for fragment ions. Identifications were stringently filtered for a FDR□<□1% at both peptide spectrum match and protein group levels. The “protein groups” MaxQuant output file was analyzed using Perseus software [55], plotting the LFQ values in volcano plots. The mass spectrometry proteomics data have been deposited to the ProteomeXchange Consortium via the PRIDE [56] partner repository with the dataset identifier PXD046528.

### RNA extraction

For RNA extraction of ribosomal fractions, each fraction was digested at 37°C for 90 min in a mix of proteinase K (final concentration 100 μg/mL; Sigma-Aldrich) and 1% SDS. Phenol acid chloroform (5:1; Sigma-Aldrich) and 10mM NaCl were then added. Samples were centrifuged at 16,000 RCF for 5 min at 4°C. The upper aqueous phase was transferred to a new tube, and 1 mL of isopropanol was added. Samples were stored at −80°C overnight to precipitate the RNA. The following day, the samples were centrifuged at 16,000 RCF for 40 min at 4°C. The supernatant was discarded, and the pellet was washed with 500 μl of 70% EtOH, then centrifuged again at 16,000 RCF for 5 min at 4°C. Pellet were air dried and resuspended in nuclease-free water. For all the other uses, RNA was extracted with TRIzol (Invitrogen) and subjected to DNase treatment using TURBO DNA-free™ Kit (Invitrogen) according to manufacturer’s instructions.

### RT and qPCR

RNA was reverse transcribed using the High-Capacity complementary DNA Reverse Transcription Kit (Thermo Fisher Scientific) on a Veriti 96-well thermal cycler (Thermo Fisher Scientific). Gene expression was measured by qPCR on a QuantStudio 5 (Thermo Fisher Scientific) and normalized with ΔΔCt method using 28S and 18S as reference genes (for polysome profiling experiments) or on the average of GAPDH, UBC, and βActin. Sequences of the primers are indicated in Table S3.

For the evaluation of lncRNA polyadenylation total RNA was reverse transcribed with the High-Capacity complementary DNA Reverse Transcription Kit (Thermo Fisher Scientific) using either random hexamers or Oligo(dT)12-18 Primer (Thermo Fisher Scientific), on a Veriti 96-well thermal cycler (Thermo Fisher Scientific). Polyadenylation was measured by qPCR on a QuantStudio 5 (Thermo Fisher Scientific) and normalized with ΔΔCt method using transcript-specific primers. GAPDH, UBC, and βActin as reference genes Sequences of the primers are indicated in Table.

### RNA sequencing and data analysis

Samples were prepared for sequencing using TruSeq Stranded Total RNA kit (Illumina) according to the manufacturer’s instructions. Libraries were sequenced on an Illumina NextSeq and HiSeq4000 according to the manufacturer’s instructions, generating PE high output configuration cycles: (R1: 100) − (I1: 6) − (I2: 0) − (R2: 100) and (R1: 50) − (I1: 6) − (I2: 0) − (R2: 0). Differential gene expression analysis was performed as previously described [23].

### Nuclear fractionation

Melanoma cells were plated in 15 cm dishes and treated with 20 μM Salubrinal (Sigma-Aldrich) for 72 h. At endpoint, cells were washed 2x with cold PBS (Gibco BRL Invitrogen), and then collected in 1,5 mL of Nuclei EZ lysis buffer (Sigma Aldrich) supplemented with 20 U/μl SUPERase-IN RNase Inhibitor (Invitrogen, Thermo Fisher Scientific), 1× Halt Protease and Phosphatase Inhibitor Single-Use Cocktail (Life Technologies) and 1 mM DTT (Sigma Aldrich). Cell lysate was thoroughly vortexed 8 times, then incubated 15 minutes on ice and centrifuged at 500 RCF for 5 minutes at 4 °C. Pellet was washed with 1,5 mL of of Nuclei EZ lysis buffer (Sigma Aldrich) supplemented with 20 U/μl SUPERase-IN RNase Inhibitor (Invitrogen, Thermo Fisher Scientific), 1× Halt Protease and Phosphatase Inhibitor Single-Use Cocktail (Life Technologies) and 1 mM DTT (Sigma Aldrich), vortexed, incubated on ice for 10 minutes on ice and centrifugated at 500 RCF for 5 minutes at 4 °C. This step was repeated once, excluding the incubation on ice. The lysate was then splitted in two for protein extraction and RNA extraction for further analysis. Nuclear enrichment was validated by RT-qPCR and by western blot.

### Protein extraction and western blot

Proteins were extracted resuspending cellular pellet in RIPA buffer [150 mM NaCl, 50 mM Tris-HCl pH 8, 1% Nonidet P40 (Thermo Fisher Scientific), 0.5% Sodium Deoxycholate (Sigma Aldrich), 1mM EDTA (Sigma Aldrich)] supplemented with 1× Halt Protease and Phosphatase Inhibitor Single-Use Cocktail (Life Technologies).

Western blotting experiments were performed using the following primary antibodies at the indicated dilution: vinculin (V9131, clone VIN-1; Sigma-Aldrich, 1:5,000), MITF (ab12039; Abcam, 1:1,000), ATF4 (#11815, clone D4B8; Cell Signaling Technology, 1:500), eIF2α-Tot (#5324, clone D7D3; Cell Signaling Technology, 1:1,000), eIF2α-Phospho-S51 (#3398, clone D9G8; Cell Signaling Technology, 1:800), ⍰-O-Linked N-Acetylglucosamine (MABS157, clone RL2; Sigma Aldrich, 1:800), ⍰-human MHC class I HLA-A, HLA-B, HLA-C (#78792101, clone W6/32; inVivoMab™, 1:1000), ⍰-human MHC class II HLA-DP/DQ/DR (#47AU4, clone IVA12; antibodies online, 1:1000), β-Actin (4970S, clone 13E5, Cell Signaling Technology, 1:3000), Recombinant anti PD-L1 (ab205921, clone 28-8, abcam, 1:1000), DAZAP1 (sc-373987, clone D-9, Santa Cruz Biotechnolgy, 1:1000), RPL11 (ab79352, clone 3A4A7, abcam, 1:1000), RPS6 (ab225676, clone EPR22168, abcam, 1:1000), RPL22 (sc-373993, clone D-7, Santa Cruz Biotechnolgy, 1:1000), RPS15 (ab157193, clone EPR11104, abcam, 1:1000).

The following HRP-linked secondary antibodies were used: ⍰-mouse IgG (NA931-1ML; Sigma-Aldrich, 1:5000) and ⍰-rabbit IgG (NA934-1ML; Sigma-Aldrich, 1:5,000). Relative protein levels were measured using ImageJ.

### Immunofluorescence

For immunofluorescence, cells were grown either on Falcon® 8-well Culture Slide (Corning) or in PhenoPlate 96-well (Perkin Elmer), fixed in 3.7% formaldehyde for 20 minutes at RT and permeabilized in 1% BSA (Sigma-Aldrich) and 0.2% Triton X-100 (Sigma-Aldrich)-containing buffer for 10 minutes on ice. Blocking was performed in 1% BSA and 10% goat serum (Abcam) for 30 minutes at RT. Primary antibody incubation was performed at RT for 1h. For co-staining of multiple antigens, sequential staining with different primary antibodies was performed. To detect MHC class I, a mouse antibody (#78792101, clone W6/32) from inVivoMab™ was used at a concentration of 1:1000. To detect MHC class II, a mouse antibody (#47AU4, clone IVA12) from antibodies online was used at a concentration of 1:1000. To detect O-Linked N-Acetylglucosamine, a mouse antibody (MABS157, clone RL2) from Sigma Aldrich was used at a concentration of 1:1000. For double staining with these two antibodies, a sequential immunostaining protocol was used. To detect CD4, an antibody already labelled with FITC (130-113-775, clone REA623) from Miltenyi biotec was used at a concentration of 1:100. To detect CD8, an antibody already labelled with PE-Vio-770 (130-113-721, clone REA734) from Miltenyi biotec was used at a concentration of 1:100. Secondary antibody incubation was performed for 45 minutes at RT in the dark. To detect calreticulin, a monoclonal rabbit antibody (Abcam #ab92516) was used at a concentration of 1:500. For RNA G4 staining, cells in culture were incubated with 1μM QUMA1 for 15 minutes in the incubator before fixation.

As secondary antibodies, ⍰-rabbit or ⍰-mouse AlexaFluor-488 and AlexaFluor-647 (Life Technologies, 1:1,000) were used. Nuclei were stained with 1:1000 DAPI (Sigma Aldrich) for 15 minutes at RT before mounting the slides with ProLong™ Glass Antifade mounting medium (Thermo Fisher Scientific). For FISH combined with immunofluorescence, cells were plated on 8-well chamber slides (Ibidi) and imaged using a Zeiss Celldiscoverer 7 with a 50x Plan Apochromat water immersion objective (NA 1.2) and an LSM900 camera with Airyscan 2 detector. A 10-z-stack per fluorophore was acquired for each image. Image processing and analysis were performed using Zeiss Zen software (blue edition) and ImageJ. Slides were imaged using Nikon C2 confocal microscope while the 96 wells were imaged by the high content screening microscope Operetta CLS+ Twister (Perkin Elmer). Analysis was performed using Image J and Harmony software (Perkin Elmer). Co-localisation analysis was performed using Fiji Image J plug-in JACoP; Van Steensel on a z-stack of confocal images. Evaluation of the level of co-localisation was measured with Cross correlation function (CCF) with a pixel shift of δ = ±20.

### Fluorescent in situ hybridization

For detection of *LISRR*, 8 FISH probes (IDT) (Table S3) marked with FAM were designed using the Stellaris probe designer software (Biosearch Technologies). Cells were grown either on Falcon® 8-well Culture Slide (Corning) or on round glass coverslips (VWR), fixed in 3.7% formaldehyde 10 minutes at RT and permeabilized in 70% EtOH for at least 1h at 4 °C. Cells were then washed twice with wash buffer (2×sodium lauryl sulfate, 10% formamide) and hybridized with a 250 nM pool of the 8 probes (for the sequences refer to table) in hybridization buffer (2×sodium lauryl sulfate, 10% formamide and 10% dextran) overnight at 37 °C. Detection and probes for rRNA FISH were performed as previously described [57].

Cells were washed with wash buffer for 30 minutes at 37 °C, twice with PBS and mounted with Invitrogen™ ProLong™ Gold Antifade mounting media with 4,6-diamidino2-phenylindole (DAPI) (Thermo Fisher Scientific). Nikon C2 confocal microscope was used for visualization and analysis was performed on Image J. Zeiss LSM980 Airyscan2 was used for the visualization of 18S, 28S and *LISRR* and co-localisation analysis was performed using plug-in JACOP on Image J. FISH images were quantified with the FISH quant Big-FISH package [58].

### TEsticular Sperm Extraction (TESE)-derived samples

Pseudonymised TESE samples were obtained from the UZ Leuven Biobank under ethical approval S67591. For TESE staining, samples were first deparaffinized in xylene and then washed in graded ethanol before permeabilization in 70% ethanol for minimum 1h at 4°C. Blocking of unspecific binding was performed using 10% goat serum in PBS for 30 minutes at RT. They were then washed with wash buffer (2×sodium lauryl sulfate, 10% formamide) and hybridized with a 250 nM pooled *LISRR* probes (for the sequences refer to Supplementary table III) and anti-DAZAP1 (A303-985A Bethyl Laboratories) 1:1000 in hybridization buffer (2×sodium lauryl sulfate, 10% formamide and 10% dextran) overnight at 37 °C. TESE samples were then washed with wash buffer for 30 minutes at 37°C before the incubation with the ⍰-rabbit AlexaFluor-647 secondary antibody (Life Technologies, 1:1000), performed for 45 minutes at RT in the dark. Nuclei were stained with 1:1000 DAPI (Sigma Aldrich) for 15 minutes at RT before mounting the slides with ProLong™ Glass Antifade mounting medium (Thermo Fisher Scientific). Samples were imaged using Nikon C2 confocal microscope.

### PDX models

The cutaneous melanoma PDX models are part of the Trace collection (https://gbiomed.kuleuven.be/english/research/50488876/54502087/Trace) and were established using left over tissue from metastatic melanoma lesions derived from patients undergoing surgery as part of standard treatment at UZ Leuven. All patients were asked to sign a written informed consent and procedures involving human samples were approved by the UZ Leuven/KU Leuven Medical Ethical Committee (S63799). PDX models Mel-006, Mel-015, and Mel-020 were derived from a female, male, and female drug-naive patients, respectively. The uveal melanoma Mel-077 sample was derived from a male patient progressing on pembrolizumab and temozolomide. Mel-018, Mel-021, and Mel-078 were derived from male, female, and male patients, respectively. PDX models Mel-015, MEL058 and MEL029 are derived from male drug-naïve patients. PDX model MEL083 is derived from a male patient resistant to targeted and immunotherapy. All the PDX models were used in accordance with the principles of the Declaration of Helsinki and with GDPR regulations. All the animal experiments were approved by the KU Leuven animal ethical committee under ECD P164/2019, P210/2018 and P032/2022 and performed in accordance with the internal, national, and European guidelines of animal care and use. Mice were implanted with tumor pieces subcutaneously in the interscapular fat pad of NSG immunocompromised mice (JAX:005557 [59]) and maintained in a semi-specific pathogen–free facility under standard housing conditions with continuous access to food and water. The health and welfare of the animals was supervised by a designated veterinarian. The KU Leuven animal facilities comply with all appropriate standards (cages, space per animal, temperature [22°C], light, humidity, food, and water), and all cages are enriched with materials that allow the animals to exert their natural behaviour. Mice used in the study were maintained on a diurnal 12-h light/dark cycle.

NOD/Scid/IL-2Rγnull, engrafted at the age of 5 weeks with CD34^+^ hematopoietic stem cells, isolated from 3 different donor cord blood, and boosted for myeloid lineage through the transient expression of human GM-CSF, IL-3, IL-4 and FLT3L, were provided by TransCure bioServices. Tumours were engrafted into humanized mice (23 weeks old) and treatment with ASO (15mg/kg sub cutaneous every second day) and/or nivolumab (10 mg/kg twice a week i.p.) was initiated when the tumor reached the size of 100 mm^3^. The study was terminated when the tumor volume reached 1.500mm^3^.

The percentage of humanization was estimated by flow cytometry analysis for CD45+ cells 14 weeks after engraftment and calculated as follows Humanization rate (%) = hCD45/(hCD45+mCD45)*100.

Mice engrafted with CD34^+^ from the same donor where randomized in the different treatment groups. According to animal welfare guidelines, mice were sacrificed when reaching humane endpoints: tumors reached a volume of 1.500 mm3 or when body weight decreased >20% from the initial weight. Mice used in this study never reached or exceeded these limits.

### ATF4 reporter

To generate the ATF4 inducible reporter, the human CARS promoter sequence (a known ATF4 target whose promoter contains a C/EBP-ATF response element [38]) was placed upstream of a luciferase-tdTomato transgene.

Specifically, the DNA sequence of the human CARS promoter (a DNA fragment of 518 bp upstream and 959 bp downstream of the human CARS TSS (ENSG00000110619, transcript ENST00000397111) was transferred from a pGL3.1 backbone (kind gift of Prof. Michael Kilberg, [39]) into a plasmid expressing both luciferase and tdTomato (pFULT, kind gift of Prof. Ghambir) using InFusion (Takara Bio) according to the manufacturer’s specifications.

### TCGA analysis

*LISRR* RNA expression levels were quantified (Log2 normalised counts per million) in 175 melanoma patients using the UCSC Xena (https://xenabrowser.net/). Patients were split in two groups, proliferative (n= 82) and invasive (n=93) according to the criteria described in [36].

*LISRR* copy number (Log2 normalised) was quantified in 473 melanoma patients (TCGA_SKCM) using the UCSC Xena (https://xenabrowser.net/).

### Bulk RNA sequencing and GSEA

Raw FASTQ files downloaded from the GEO database [60] project “mRNA expressions in pre-treatment melanomas undergoing ⍰-PD-1 checkpoint inhibition therapy” (data accessible at NCBI GEO database [34], accession GSE78220) or produced by the laboratory were passed through FASTQ Groomer [61]. Adaptors were trimmed and low-quality reads were discarded using Trimmomatic [62]. The reads were pseudo-aligned to the human reference genome GRCh38 [63] and counted using Kallisto [64]. The R package Sleuth [65] was used for normalization of the counts. was handled with the R package Seurat. The R packages ggpubr and ggplot were used to plot gene expression. Geneset enrichment analysis [66] was performed with GSEA 4.1.0 on the normalized counts. Plots visualizing GSEA data were made in R using the ggplot2 package [67] and only genesets with nominal p-value < 0.05 and FDR q-value < 0.25 are shown.

### Single cell RNA sequencing analysis

The single cell sequencing data from human melanoma patients before and after a single dose of ICB therapy [36] was handled with the R package Seurat. The R packages ggpubr and ggplot [68] were used to plot gene expression from this dataset.

### Molecular Cartography

#### Probe Design

The probes for 100 genes (table III) were designed at the gene-level using Resolve’s proprietary design algorithm. For every targeted gene all full-length transcript sequences from the ENSEMBL database were used as design targets if the isoform had the GENCODE annotation tag ‘basic’ [69, 70]. To filter highly repetitive regions, the abundance of k-mers was obtained from the background transcriptome using Jellyfish [71]. Every target sequence was scanned once for all k-mers, and those regions with rare k-mers were preferred as seeds for full probe design. A probe candidate was generated by extending a seed sequence until a certain target stability was reached. A set of simple rules was applied to discard sequences that were found experimentally to cause problems. After these fast screens, every kept probe candidate was mapped to the background transcriptome using ThermonucleotideBLAST [72] and probes with stable off-target hits were discarded. Specific probes were then scored based on the number of on-target matches (isoforms), which were weighted by their associated APPRIS level [73], favouring principal isoforms over others. From the pool of accepted probes, the final set was composed by greedily picking the highest scoring probes. Gene and probes ID are provided in Supplementary Table 3.

#### Sample preparation

Freshly dissected humanized PDX tumours (3 per treatment cohort) were frozen in isopentyne (Fisher Scientific) and sectioned with a cryostat HM 525 NX (Thermo Fisher) and 10 µm-thick sections were placed within the capture areas of cold Resolve Biosciences slides. Samples were then sent to Resolve BioSciences on dry ice for analysis. Upon arrival, mouse tissue sections were thawed and fixed with 4% v/v Formaldehyde (Sigma-Aldrich F8775) in 1x PBS for 20 minutes at 4 °C, whereas the human samples were fixed for 45 minutes at 4 °C. After fixation, sections were washed twice in 1x PBS for two min, followed by one min washes in 50% ethanol and 70% ethanol at RT. Fixed samples were used for Molecular Cartography^TM^ (100-plex combinatorial single molecule fluorescence in-situ hybridization) according to the manufacturer’s instructions as previously described [74]. A total of 50 tiles per sample were imaged.

#### Imaging

Samples were imaged on a Zeiss Celldiscoverer 7, using the 50x Plan Apochromat water immersion objective with an NA of 1.2 and the 0.5x magnification changer, resulting in a 25x final magnification. Standard CD7 LED excitation light source, filters, and dichroic mirrors were used together with customized emission filters optimized for detecting specific signals. Excitation time per image was 1000 ms for each channel (DAPI was 20 ms). A z-stack was taken at each region with a distance per z-slice according to the Nyquist-Shannon sampling theorem. The custom CD7 CMOS camera (Zeiss Axiocam Mono 712, 3.45 µm pixel size) was used. For each region, a z-stack per fluorescent colour (two colours) was imaged per imaging round. A total of 8 imaging rounds were done for each position, resulting in 16 z-stacks per region. The completely automated imaging process per round (including water immersion generation and precise relocation of regions to image in all three dimensions) was realized by a custom Python script using the scripting API of the Zeiss ZEN software (Open application development).

#### Spot Segmentation

The algorithms for spot segmentation were written in Java and are based on the ImageJ library functionalities. Only the iterative closest point algorithm is written in C++ based on the libpointmatcher library (https://github.com/ethz-asl/libpointmatcher).

#### Pre-processing

As a first step all images were corrected for background fluorescence. A target value for the allowed number of maxima was determined based upon the area of the slice in µm² multiplied by the factor 0.5. This factor was empirically optimized. The brightest maxima per plane were determined, based upon an empirically optimized threshold. The number and location of the respective maxima was stored. This procedure was done for every image slice independently. Maxima that did not have a neighboring maximum in an adjacent slice (called z-group) were excluded. The resulting maxima list was further filtered in an iterative loop by adjusting the allowed thresholds for (Babs-Bback) and (Bperi-Bback) to reach a feature target value (Babs: absolute brightness, Bback: local background, Bperi: background of periphery within 1 pixel). This feature target values were based upon the volume of the 3D-image. Only maxima still in a zgroup of at least 2 after filtering were passing the filter step. Each z-group was counted as one hit. The members of the z-groups with the highest absolute brightness were used as features and written to a file. They resemble a 3D-point cloud. Final signal segmentation and decoding: To align the raw data images from different imaging rounds, images had to be corrected. To do so the extracted feature point clouds were used to find the transformation matrices. For this purpose, an iterative closest point cloud algorithm was used to minimise the error between two point-clouds. The point clouds of each round were aligned to the point cloud of round one (reference point cloud). The corresponding point clouds were stored for downstream processes. Based upon the transformation matrices the corresponding images were processed by a rigid transformation using trilinear interpolation. The aligned images were used to create a profile for each pixel consisting of 16 values (16 images from two colour channels in 8 imaging rounds). The pixel profiles were filtered for variance from zero normalized by total brightness of all pixels in the profile. Matched pixel profiles with the highest score were assigned as an ID to the pixel. Pixels with neighbours having the same ID were grouped. The pixel groups were filtered by group size, number of direct adjacent pixels in group, number of dimensions with size of two pixels. The local 3D-maxima of the groups were determined as potential final transcript locations. Maxima were filtered by number of maxima in the raw data images where a maximum was expected. Remaining maxima were further evaluated by the fit to the corresponding code. The remaining maxima were written to the results file and considered to resemble transcripts of the corresponding gene. The ratio of signals matching to codes used in the experiment and signals matching to codes not used in the experiment were used as estimation for specificity (false positives).

#### Downstream analysis

For the analysis, cell segmentation was performed with the Automatic Cell Segmentation tool provided by Resolve based on DAPI images. Pixels are burred at tiling gaps in the DAPI panorama images to migitate over-segmentation due to artificial borders at the edges of field of views with costum software called MindaGap at default settings MindaGap is available at https://github.com/ViriatoII/MindaGap. Subsequently, Cellpose [75] was used to segment the cell nuclei in the pre-trained cyto model with diameter and flow-thresh parameters set to 50.0 and 0.5, respectively. Compartment specific genes were determined by calculating the ratio of total transcript counts and transcripts inside DAPI segments per gene. Genes having >=70% of transcripts located inside DAPI segments are considered nucleus specific, genes with <=30% of transcripts in DAPI segments are considered cytoplasmic. Cells were segmented based on transcript coordinates using Baysor [76] with DAPI segments as prior and the following parameter settings: --no-ncv-estimation --force- 2d --n-clusters=1 --prior-segmentation-confidence 0.9 -m 3. Baysor’s compartment genes feature was used by adding genes matching the above-mentioned compartment definitions to Baysor’s --nuclei-genes and --cyto-genes parameters, respectively. Finally, the outline of cells was predicted by Baysor by running the convex hull algorithm on the corresponding subset of transcripts assigned to each individual cell. The R package Seurat was used to normalize and integrate the count matrices into one dataset. Cells that expressed less than two unique genes and had less than 10 counts have been filtered out. Gene expression of different immune cell and state markers (Supplementary table 3) and lncRNAs were plotted with the R packages ggpubr and ggplot [68]. Differentially expressed genes were identified with the pseudobulk aggregation by summing method [77].

#### WGA staining

Existing solution was discarded and HBSS was used to wash the slides twice for 1min. The solution was aspirated and discarded. Glycans were stained with Wheat Germ Agglutinin, Alexa Fluor™ 633 Conjugate as previously described [78].

### CCLE

*LISRR* Transcripts Per Million (TPM) counts were plotted for cutaneous melanoma cell lines from the CCLE [79] Sanger GDSC1 dataset [80] and correlated (Pearson correlation coefficient) with AUC of BRAF and MEK inhibitors dabrafenib and trametinib.

### CPC2

Illumina reads were passed through FASTQ Groomer [60]. Adaptors were trimmed and low-quality reads were discarded using Trimmomatic [62]. The reads were pseudo-aligned to the human reference genome GRCh38 [63] and counted using Kallisto [64]. The R package Sleuth [65] was used for differential transcript expression analysis. The coding potential of the differentially expressed transcripts (FDR<0.05 & average control on average salubrinal <0.5 or >2) were predicted by CPC2 [81]. The R package ggplot2 [68] was used for making the figures.

### Patient derived tumour fragments (PDTF)

Fresh tumour samples from patients were collected at UZ Leuven under protocol S57760. The patient underwent a biopsy of a metastatic skin lesion that progressed on anti-PD1 based therapy. The tumour fragments were washed twice in PBS supplemented with 1% penicillin/streptomycin. The procedure of cutting and embedding in extracellular matrix was performed as previously described [81]. The day after embedding, the samples were treated with 10 µg/mL of α-PD1 (Nivolumab, Bristol-Myer Squibb) and transfected with 50 nM *siLISRR* (Sigma, for list of sequences you can check the table S1) or siCtrl (MISSION® siRNA Universal Negative Control #1 and #2, Sigma Aldrich). 48h after the beginning of the treatment, the PDTF were extracted from the extracellular matrix, washed with PBS and fixed 20 minutes in 4% PFA (Sigma). They were then washed and stored in 70% EtOH for later embedding in Paraffin and Akoya/Opal staining [74].

### Search for RNA binding protein consensus

Genes depleted and enriched at the polysomes upon *LISRR* knockdown as determined by RIVET were uploaded into the oRNAment database to search for binding sites of RNA binding proteins. A cutoff score of 1 was used for binding site inclusion in the analysis.

## Supporting information

Mass Spectrometry

Proteomics on ribosomal fractions

Resolve probes

## Author’s contributions

SC performed most of the in vitro wet lab experiments and assembled figures. YV performed all the bioinformatic analysis. ACC provided support for bulk RNAseq analysis. AM performed RNA pull-downs and RPL26 RIP. VK performed the experiments with pFULT-hCARS reporter. ZK performed experiments in supplementary figure 2CD and occasionally western blots. RV designed and built the pFULT-hCARS reporter. ED performed PDX experiments. MFB coordinated the in vivo work. EG provided ASO. OB provided access to essential human clinical samples and data. SA prepared samples for molecular cartography and stainings and quantified them. JP provided access to essential human single-cell RNAseq data and supervision for figure 6E preparation. MB provided TESE samples for DAZAP1 and *LISRR* co-localisation. ST provided analysis of mouse humanization. SH performed the experiments on MM0165. TB and AC designed, performed and interpreted mass spectrometry experiments. EL designed the study, interpreted the data and wrote the manuscript. All authors read and edited the manuscript.

## Acknowledgment

We would like to thank the Light Microscopy and Imaging Network - LiMoNe (VIB-KULeuven), especially Axelle Kerstens, Nicolas Peredo and Abril Escamilla Ayala, for the excellent technical support. We would also like to thank Dr K. De Keersmaecker and Dr J-C Marine for critical reading of the manuscript. We would like to thank Guy Schepers and Frédérick Coosemans (Rega Institute) and Lukas Vanwynsberghe (LMCB-VIB) for the excellent technical support with ASO synthesis and with AKOYA staining respectively. We would like to thank also Lucas Ferreira Maciel (LMCB-CCB VIB) for the assistance with the Resolve data representation and Chiara Ghirardi (IEO) for the technical support in sample processing prior to LC-MS/MS analysis. We would like to apologise for not being able to cite all the literature about other transcripts generated by the locus encoding for *LISRR* and not related in function nor in transcript structure to *LISRR*. The present study was funded by a KU Leuven C1 grant, a Belgian federation for Cancer grant (FAF-F/2018/1184). E. Leucci was funded by the Melanoma Research Alliance Amanda and Jonathan Eilian young investigator award. Trace staff was self-supported. T. Bonaldi is supported by the Italian Association for Cancer Research grant number IG-2018-21834. This work has been supported by EPIC-XS, project number 823839, funded by the Horizon 2020 programme of the European Union. V. Katopodi was a recipient of a FWO – Flanders research organization PhD fellowship (1S47519N). Y. Verheyden is FWO – Flanders research organization PhD fellowship (1SC5122N). S. Cinque was a recipient of FWO – Flanders research organization PhD fellowship (1SD1620N).

