## Supplementary Figures for "The assembly of cancer-specific ribosomes by the lncRNA *LISRR* suppresses melanoma anti-tumour immunity"

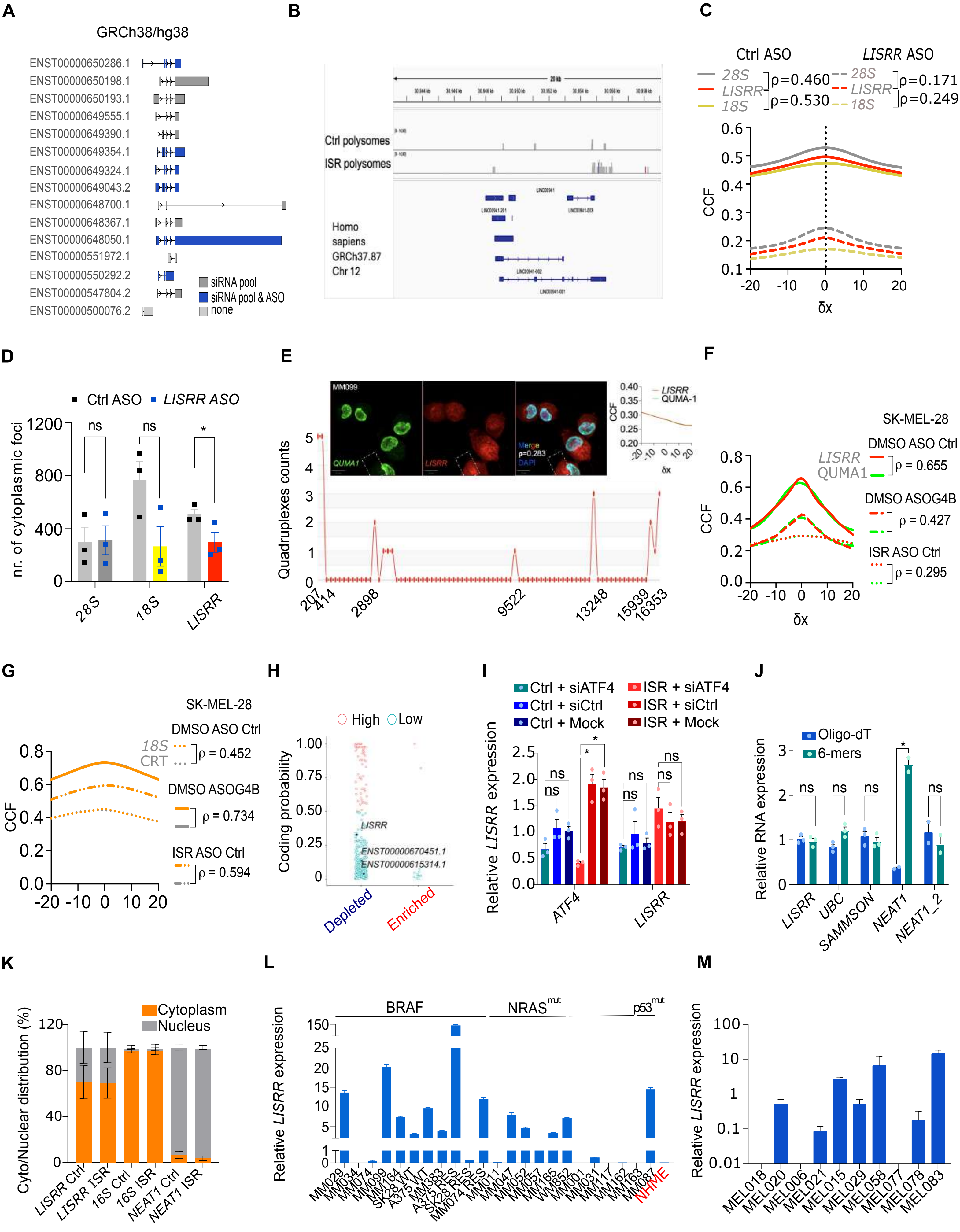


**Supplementary Figure 1A-M: LISRR is a cytoplasmic polyadenylated lncRNA**

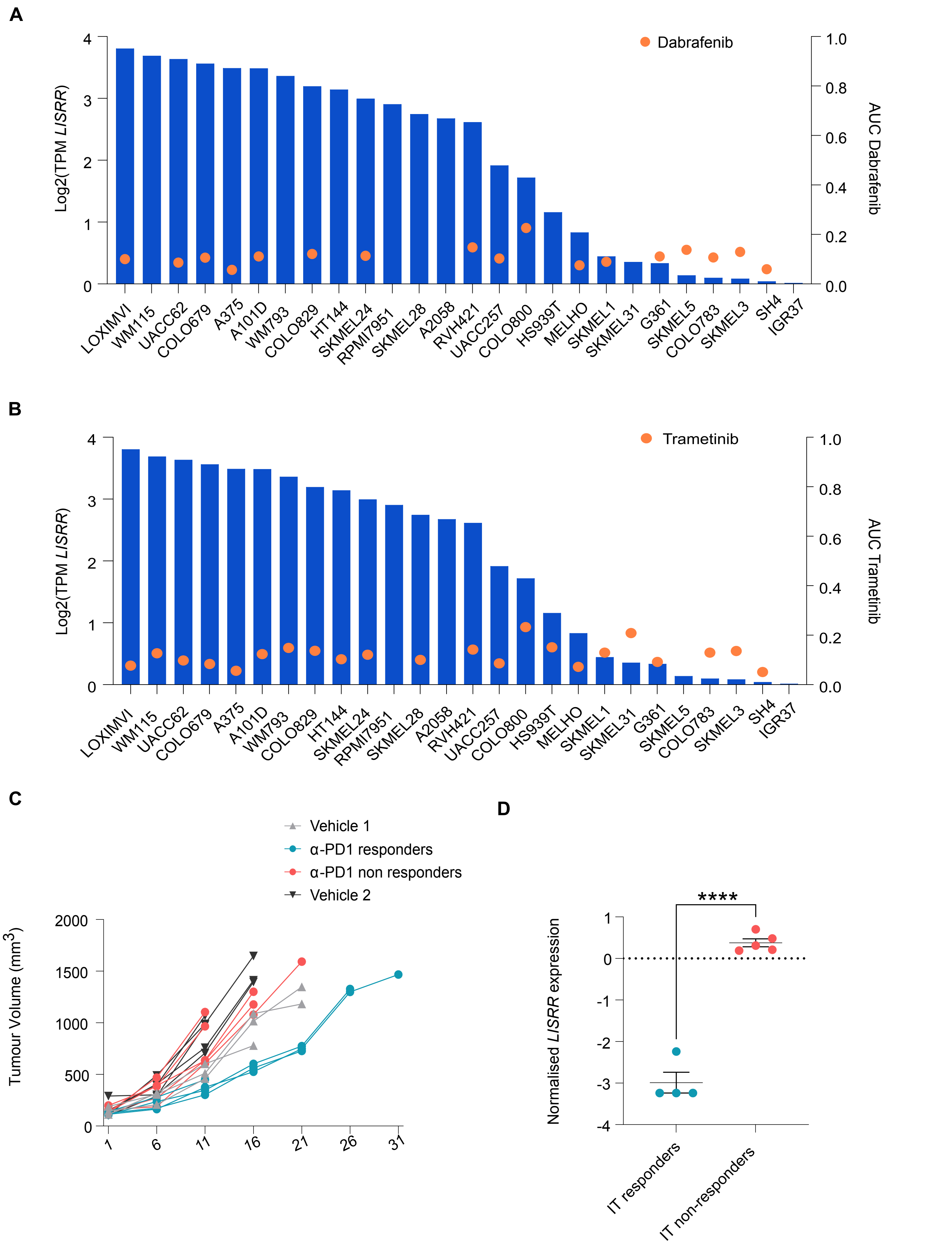


**Supplementary Figure 2A-D: LISRR expression correlates with responses to ICB in humanised PDX**

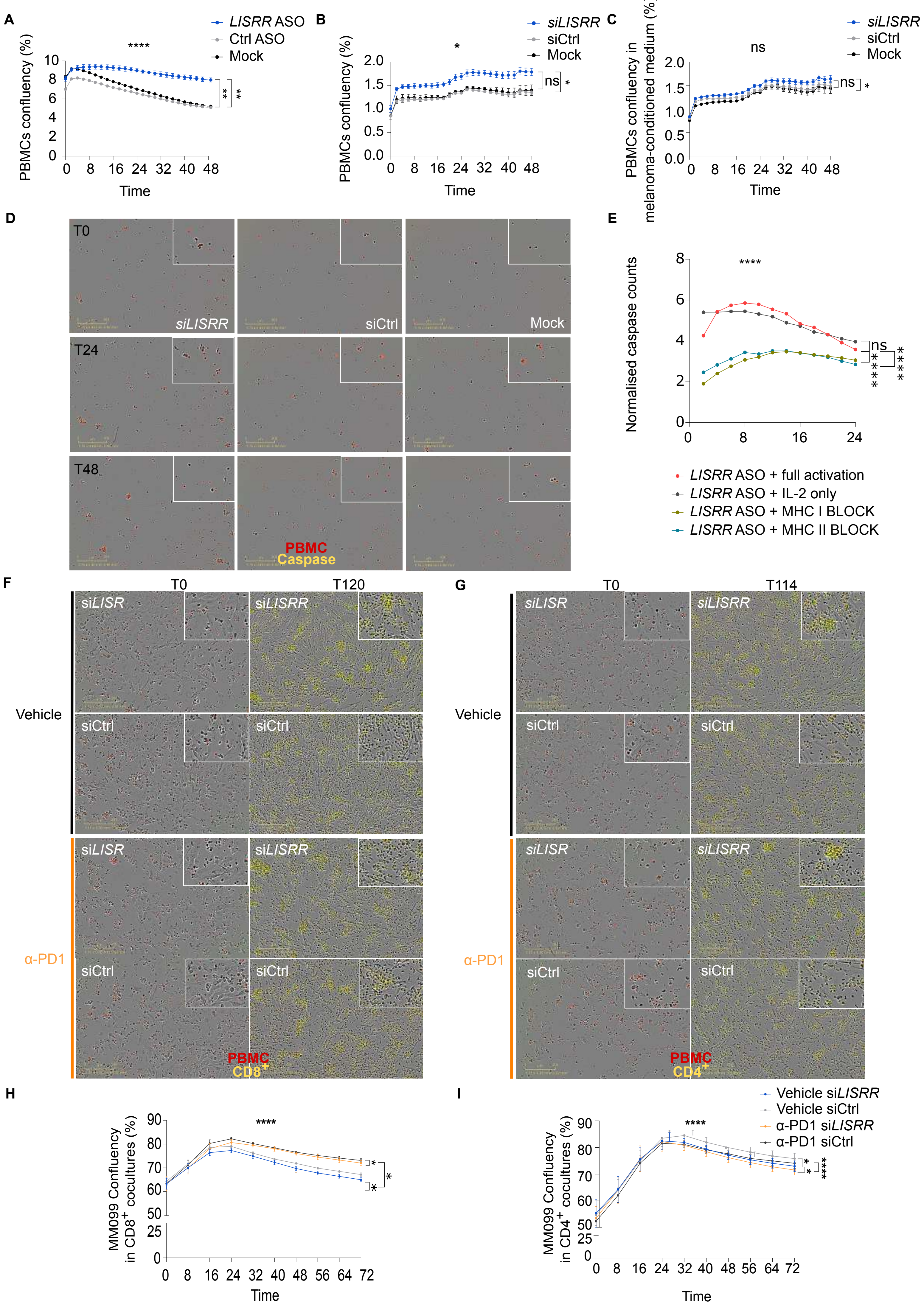


**Supplementary Figure 3A-I: Inhibition of LISRR enhances anti-melanoma immune responses**

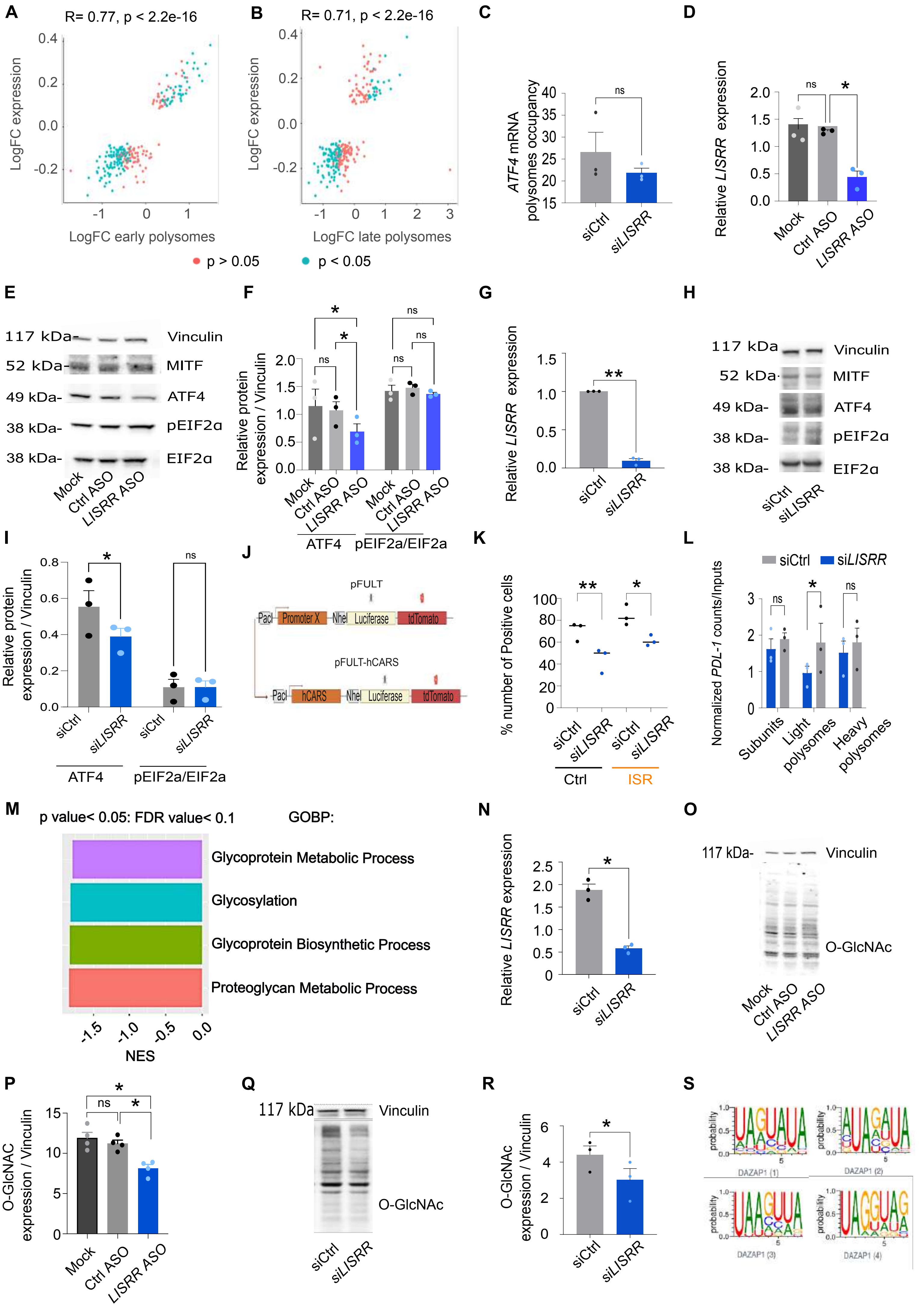


**Supplementary figure 4 A-S: LISRR regulates translation**

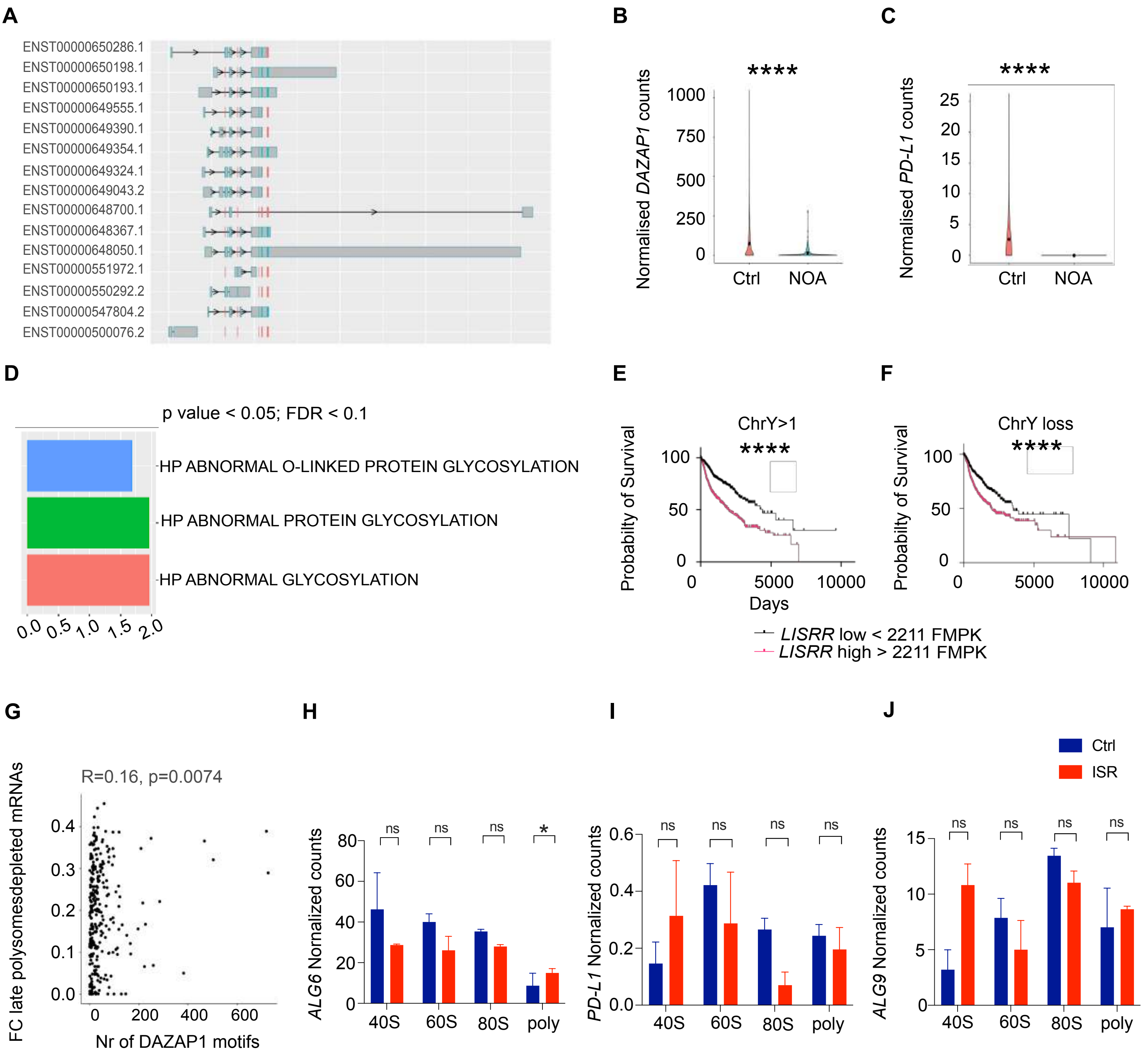


**Supplementary Figure 5A-J: DAZAP1 partly mediates LISRR biological functions**

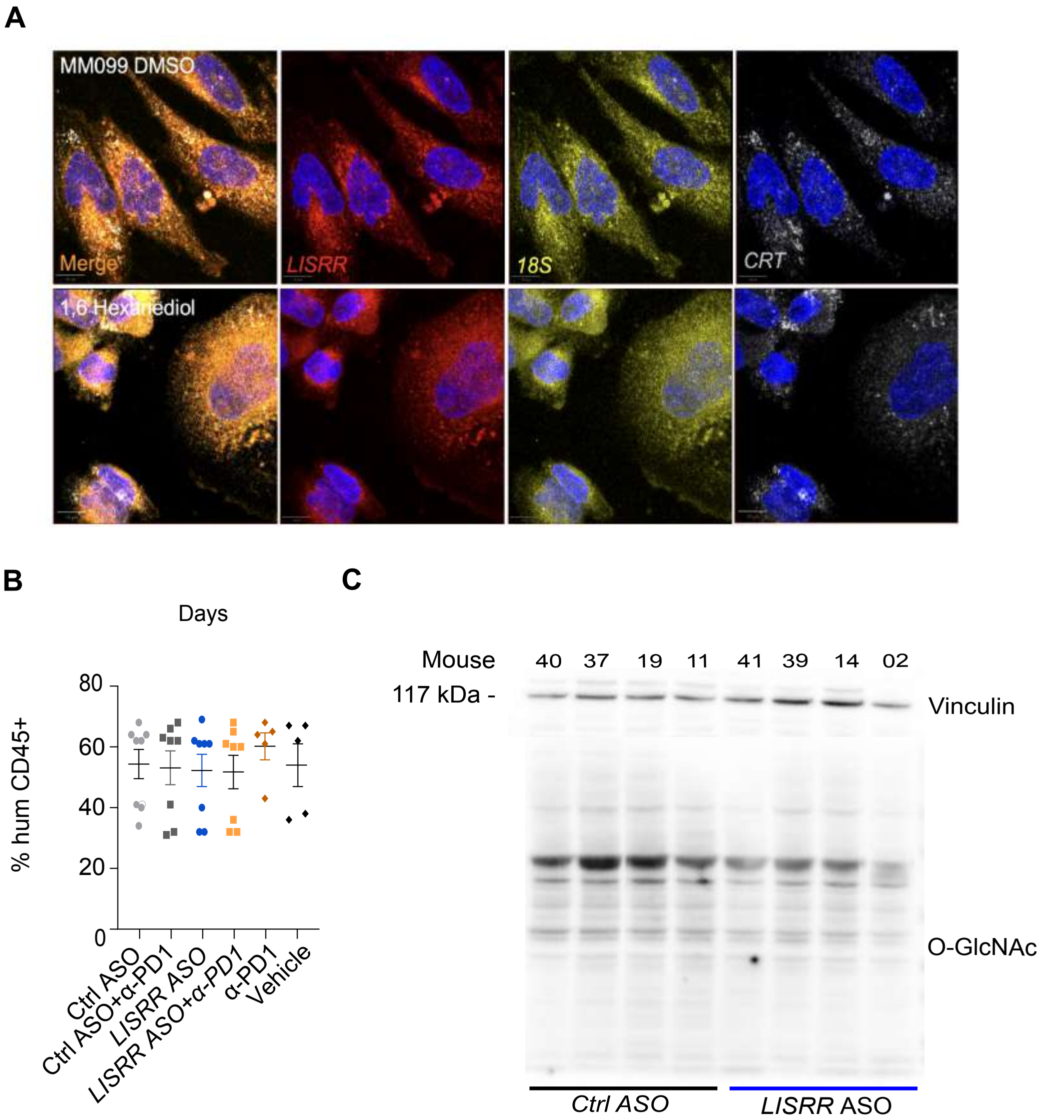


**Supplementary Figure 6A-D: Inhibition of LISR in vivo in humanised PDX models regulates glycosylation.**
